# Generation of effector CD4+ T cells from human iPSC

**DOI:** 10.1101/2025.08.01.667959

**Authors:** Julian Amirault, Dar Heinze, Mengwei Yang, Charles M. Kerr, Pushpinder S. Bawa, Laura Polanco, Gabriel M. Sgambettera, Feiya Wang, Anna C. Belkina, Jennifer E. Snyder-Cappione, Gustavo Mostoslavsky

**Affiliations:** Center for Regenerative Medicine of Boston University and Boston Medical Center; Boston, MA, USA; Department of Virology, Immunology, and Microbiology; Boston University Chobanian and Avedisian School of Medicine, Boston, MA, USA; Department of Pathology and Laboratory Medicine; Boston University Chobanian and Avedisian School of Medicine, Boston, MA, USA; Flow Cytometry Core Facility, Boston University Chobanian and Avedisian School of Medicine, Boston, MA, USA; Department of Medicine, Section of Gastroenterology at Boston University and Boston Medical Center; Boston, MA, USA

## Abstract

Off the shelf CD4+ T cell therapies, particularly those with immunoregulatory or cell repair functions, could be transformative in CAR therapies for cancers and treatment of chronic inflammatory diseases. However, progress is stunted in this area due to challenges generating human CD4+ T cells from induced pluripotent stem cells. Here we describe a key role for the withdrawal of Notch ligand during the final step of stimulation through the T cell receptor to prompt T cell maturation allowing access to the CD4 lineage in iPSC T cells (iCD4+ T cells). Functional analyses of iCD4+ T cells using a novel high-parameter CyTOF intracellular cytokine staining panel revealed both canonical Th1 cytokine signatures and cells producing varying combinations of other cytokines including IL-4, IL-8, and IL-13. Single cell RNA sequencing of iCD4+ T cells demonstrated a transcriptional signature similar to human blood CD4+ T cells. We believe this robust yet simple platform represents a key step towards the generation of off the shelf iCD4+ T cell therapies with utility for the treatment of a panoply of diseases including cancer and inflammatory autoimmune disorders.

**HIGHLIGHTS:** T cell receptor stimulation of iPSC CD4+/CD8+ T cell progenitors on coating without Notch ligand allows access to the CD4+ T cell lineage iPSC derived CD4+ T cells express varied cytokines in response to stimulation

## INTRODUCTION

T cells begin as hematopoietic progenitor cells which migrate from the bone marrow to the thymus^1,2^. Transition from hematopoietic progenitors to a CD4/CD8 double negative followed by a CD4/CD8 double positive (DP) lymphoid population takes place in the Notch ligand-rich environment of the thymic cortex, where the cells are maintained by survival cytokine IL-7^3,4^. DP progenitor cells are signaled to mature into single positive (SP) thymocytes by T cell receptor (TCR) binding to major histocompatibility complexes (MHCs) presented by thymic epithelial cells in a process termed “positive selection”^5–9^. SP semi-mature T cells undergo negative selection, where highly self-reactive clones undergo apoptosis, with the exception of some strongly signaled CD4+ clones, which are diverted to an anti-inflammatory regulatory T cell fate^7, 10^.

Despite work implicating strength and duration of TCR signaling alongside co-receptor or cytokine signals in functional lineage differentiation and commitment, the mechanisms resulting in the divergence of these two fundamental T cell lineages, CD8+ vs CD4+, remain in question^10–14^. IL-7 is an established requirement for CD8 SP T cell survival, and both CD4 and CD8 SP cells require the binding of the TCR to MHC (MHC II and MHC I respectively) loaded with a compatible antigen for lineage commitment^15^.

T cells are the basis of multiple promising cell therapies, from Chimeric Antigen Receptor (CAR) T cell approaches which target cancers to T regulatory cell and CAR T regulatory anti-inflammatory treatments for inflammatory conditions. Conventional CAR T immunotherapy approaches use either autologous T cells derived from the patient being treated or allogeneic T cells taken from healthy donors^16^. A key drawback of autologous CAR T therapy is time and cost to treatment, while donor-to-donor variation in starting T cell material can result in differences in outcome for patients. The ratio of CD4+ to CD8+ T cells alongside the presence or absence of specific T cell subpopulations have been connected to the effectiveness of treatment response in CAR T therapy^17–19^. While allogeneic CAR T can be easily prepared and banked prior, it presents significant risks of immune rejection; groups are currently working on universal donor edits to cell preparations reducing immunogenicity^16^. An alternative cell source with lower cost and rapid production when compared to autologous or allogeneic sources could rely on the generation of T cells from induced Pluripotent Stem Cells (iPSC)^20–23^. The same hypoimmunogenicity edits are easily applicable to iPSC derived T cells, enabling a much more cost efficient universal off-the-shelf therapeutic product.

While robust generation of the CD4+/CD8+ double positive (DP) progenitor cells from iPSCs have been demonstrated, generally only CD8+ cytotoxic lymphocytes have been described^22,24,25^. CD4+ T cell cytokines are important for many effective T cell therapies and CD4+ T cell populations have been shown to be highly relevant in current CAR T immunotherapies, providing important regulatory support to CD8+ T cells^26–30^. In this report, we describe a highly scalable, feeder-free platform for the generation of functional iPSC-derived CD4+ T cells, detailing their stimulation-specific combinational expression of 18 cytokines and cytotoxic effector molecules at single cell resolution using CyTOF alongside transcriptional signatures using single cell RNA sequencing.

## RESULTS

### Notch withdrawal allows for CD4+ lineage emergence

We have previously published a T cell generation protocol using a doxycycline-inducible Notch 1 intracellular domain in the first three days of hematopoietic stem and progenitor induction that robustly increases the proportion of cells which can access the T cell lineage^22^. This 12-day hematopoietic stem and progenitor protocol allowed robust generation of CD34+ progenitors (Fig. 1B, left panel). These cells are fed forward onto plates coated with Notch ligand, and after a further 28-33 days large numbers of DP progenitor cells are generated (Figure 1B, right panel). The cells express hallmark lymphoid progenitor markers, such as CD7 (Fig. 1B, center panel) at day 26 after T cell specification followed by CD3, T cell receptor, CD4 and CD8 (Fig. 1B, right panels) by day 40. Many groups, including ours, have used a combination of anti-CD3/CD28 in the presence of Notch ligand with IL-7 or IL-15 during maturation of DP to CD8 SP T cells^22,31,32^. We screened for signals that could allow or block access to the CD4+ lineage by systematically removing individual factors from the DP to SP maturation culture conditions. To our surprise, when we moved the double positive cells to RetroNectin^®^ coated wells, a fibronectin derivative lacking Notch ligand, and media supplemented with IL-7 and anti-CD3/CD28, a population of CD4 SP cells emerged. albeit with poor viability (Fig. S1). We hypothesized that the anti-CD3/CD28 stimulation was too strong resulting in negative selection and apoptosis. Therefore, we removed the anti-CD28 co-stimulatory signal; indeed, stimulation of the cells with anti-CD3 antibody alone dramatically increased viability of iT cells after the DP to SP maturation while robustly differentiating into the CD4+ T cell lineage (Fig. 1C) (Fig. S1). CD3 and TCRɑβ appeared downregulated in response to stimulation at day 54 (Fig. 1C).

**Fig. 1.**
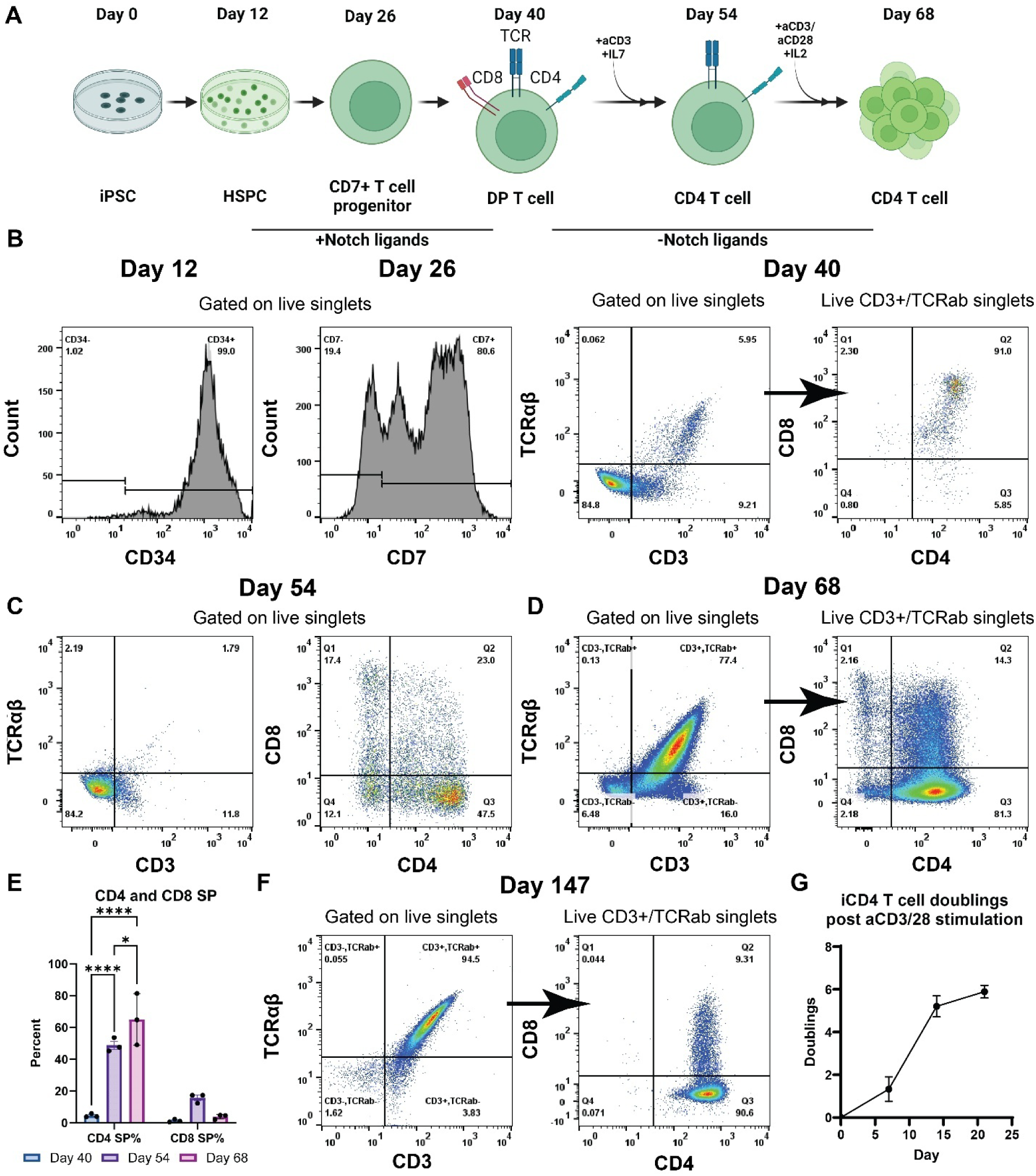
Analysis of cells from throughout the process of generating and maintaining iCD4+ T cells. (**A**) Schematic of iT differentiation to CD4+ cells. (**B**) Representative flow cytometry of key developmental timepoints and metrics of DP T cells. iPSCs were plated at day -2 and cultured for 12 days in HSPC differentiation conditions. The resultant CD34+ HSPCs were purified using magnetic beads. Day 12 HSPCs were moved to Notch-stimulatory lymphoid coating and cultured with StemCell Technologies lymphoid expansion media until day 26, where cells were assayed for CD7 expression. Day 26 cells were replated on fresh Notch stimulatory coating and cultured with StemCell Technologies lymphoid maturation media to day 40, where cells are assayed for TCR, CD3, CD4 and CD8 expression. (N=3) (**C**) Representative flow cytometry of D54 T cells, with parent gating in Figure S3A. Day 40 iPSC DP T cells were moved to Retronectin coating and treated with anti-CD3 antibody (UCHT1, 5 ug/ml), half media changes were performed every 3 or 4 days. (N=3) (**D**) Representative flow cytometry of D68 T cells post expansion. Day 54 iCD4+ T cells were moved to fresh Retronectin coated wells and treated with media supplemented with anti-CD3/CD28 and 200 U/ml IL-2. (N=3 differentiations) (**E**) Percentage of CD4 and CD8 positive T cells from iCD4+ T cell differentiation by flow cytometry at day 40, 54 and 68, N=3, analyzed by 2-way ANOVA followed by a Tukey’s multiple comparison test, *P < 0.05; **P < 0.01; ***P < 0.001; ****P < 0.0001. (**F**) Representative flow cytometry of T cells from long-term culture in X-Vivo 15 media supplemented with 200 U/ml IL-2. N=3 differentiations, parent gating for each plot as indicated. **(G)** Number of doublings of iCD4+ T cells after stimulation with anti-CD3/CD28 (N = 3 differentiations).

DP to SP specification in the thymus is correlated with loss of apoptotic responses to strong TCR stimulation^14,33^. To test the capacity of our iCD4+ cells to survive strong TCR stimulation, we treated newly specified iCD4+ cells with anti-CD3/CD28 in the presence of high-dose IL-2 (200 U/mL). CD4 single positivity was maintained throughout this expansion stimulation along with recovery of TCRɑβ and CD3 expression at day 68 (Fig. 1D, E). We found that our iPSC-derived CD4+ T cells (iCD4+ T cells) can be expanded extensively in vitro while maintaining their T cell identity for at least 147 days, following two rounds of stimulation and expansion (Fig. 1F, G). Treatment with anti-CD3/CD28 in serum-free media with 200 U/mL IL2 resulted in approximately six doublings (Fig. 1G), in line with previously reported T cell proliferation rates in the same media^34^. Importantly, we have found that iT cells can be frozen and later thawed at days 12, 26 or any time after day 68 allowing for cell banking.

### iCD4+ T cells map to thymocyte developmental stages

In order to further understand the developmental trajectory of our iCD4+ T cells, we used multi-parameter flow cytometry covering 23 developmental and functional markers (Table S1). Following the differentiation protocol outlined in Figure 1A we analyzed developing iT cells at day 40, 54, 68 and 75 of differentiation. As expected, most of the DP cells at day 40 were negative for or weakly expressed many of the maturation markers, including CD5, CCR7, CD69, MHC I and CD25 (Fig. 2).

**Fig. 2.**
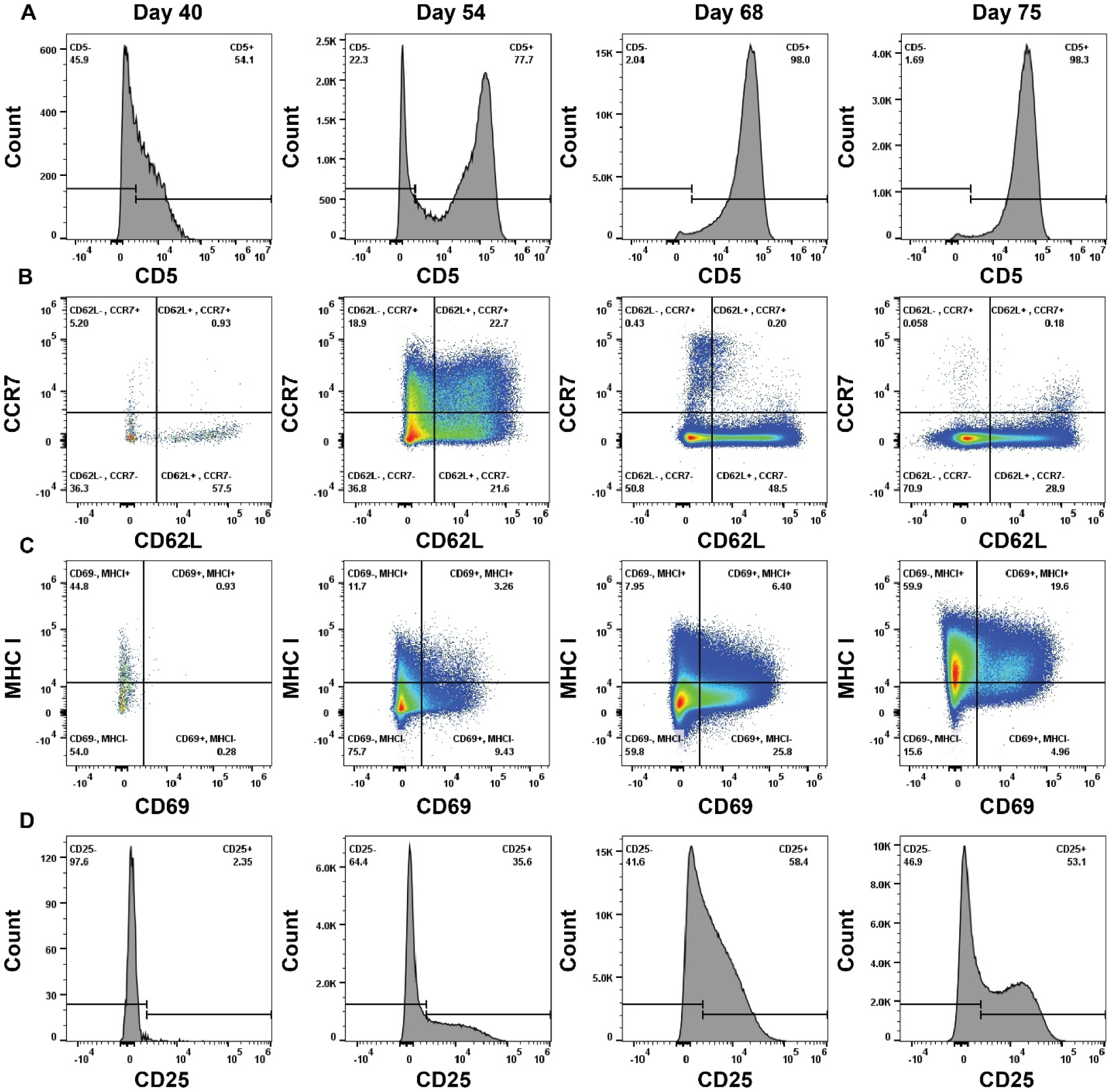
Maturation marker flow cytometry of multiple time points of CD4 specification (day 40 and day 54) and expansion (day 68 and day 75) of iT cells. The populations shown are gated on Live/Single/CD45+/CD14-/CD16-/CD56-/CD335-/CD337-/CD3+/ TCRɑβ+ cells, with the exception of Day 54, which is gated on Live/Single/CD45+/CD14-/CD16-/CD56-/CD335-/CD337- cells due to poor CD3 and TCRɑβ expression. Parent gating shown in figure S3B, representative of N=3 differentiations. iPSC DP T cells were specified to iCD4+ T cells using the differentiation outlined in fig. 1A and analyzed for expression of CD5 (**A**), CD62L and CCR7 (**B**), CD69 and MHC I (**C**), and CD25 **(D)**.

Maturation of the cells induced strong upregulation of CD5 throughout the course of CD4 specification, with a substantial population of CD5hi cells emerging by day 54 while nearly all cells become CD5hi by day 68 and day 75 (Fig. 2A). CD5 intensity has been shown to be reflective of TCR stimulation during selection in primary thymocytes^35,36^.

CCR7 is upregulated during primary T cell development within positive selection and directs cells to the thymic medulla^37,38^. We observed upregulation of CCR7 on our iT cells at day 54 of culture corresponding to a post-selection timepoint. CCR7 expression decreases by day 68 and 75, likely reflecting restimulation of the TCR and CD28 triggering further specification toward an effector phenotype (Fig 2B). CD62L (L-Selectin) has been reported to be upregulated in the most mature SP thymocytes as well as mature naive T cells ^39^. In our system, CD62L is expressed early on and became highly expressed in a population of the SP cells upon maturation at day 54. Unlike CCR7 however, CD62L expression is maintained through day 75 in a substantial population of the iCD4+ T cells (Fig 2B).

Further evidence for positive selection in our iCD4+ T cells can be provided through CD69 and MHC class I expression ^40^. CD69- DP cells represent a pre-selection group of DP T cells, while CD69+ cells appear in the lineage commitment process post-selection. CD69, while likely upregulated following the second stimulation of the SP T cells at day 54, is downregulated by day 75 of differentiation (Fig. 2C). Similarly MHC-I, initially not expressed in DP cells, becomes highly expressed by day 75 and its expression correlated well with the acquisition of proliferative capability by the SP iT cells (Fig. 2C)^14^. CD25 expression, which is negative at day 40, increases throughout the differentiation (Fig. 2D). CD25 expression likely reflects activation and indicates capacity for IL-2 responsiveness in iT CD4+ cells, as has been shown in primary effector CD4+ T cells^41^.

In order to definitively confirm the identity of our iCD4+ T cells we analyzed expression of the transcription factor ThPOK (also known as *ZBTB7B*), a key CD4+ T cell transcription factor ^42^. Transient upregulation of ThPOK in DP T cells transitions into stable expression in mature CD4+ T cells^13^. As detected by RT-qPCR, iCD4+ T cells clearly express ThPOK at every timepoint albeit at lower expression levels compared to PBMC (Fig. S2C). Importantly, iCD4+ T cells possess diverse TCRβ V and J usage, indicative of multiclonal iCD4+ T generation (Fig. S4), further evidence that our protocol recapitulates normal T cell development.

### Cytokine analysis of iCD4+ T cells at single cell resolution reveals hallmark Th1 cells and other T helper expression signatures

Next, we sought to determine the phenotypic and functional signatures of the day 68 iCD4+ T cells using a novel 30+ metal intracellular cytokine CyTOF panel (Table S2). After short-term (overnight) culture with either media alone or with phorbol 12-myristate 13-acetate (PMA) and ionomycin, iCD4+ T cells were stained and run on the CyTOF. T cell lineage markers showed all cells were T cells (CD3+), with >90% CD4 single positive, and a negligible percentage of cells expressing CD8 (Fig. 3A, S5). Collected data was visualized into 2D space using opt-SNE and clustered via Phenograph from expression of both surface and intracellular markers^43,44^. The vast majority of iCD4+ T cells produced both IFN-γ and TNF-α (Fig. 3B, C); this finding aligned with analysis of CD4+ T cells from PBMC (Fig. 3D) and CAR-T populations^45,46^. Importantly, inclusion of several more cytokine readouts revealed diversity of effector function profiles within this ‘Th1’ iCD4+ T cell population, including cells producing both Th2 cytokines IL-4 and IL-13, a notable proportion producing IL-8, co-expression of MIP-1α, and MIP-1β, and perhaps most strikingly, five clusters containing strong producers of the cell repair cytokine amphiregulin in a stimulation-specific manner (Fig. 3B, C, D).

**Fig. 3.**
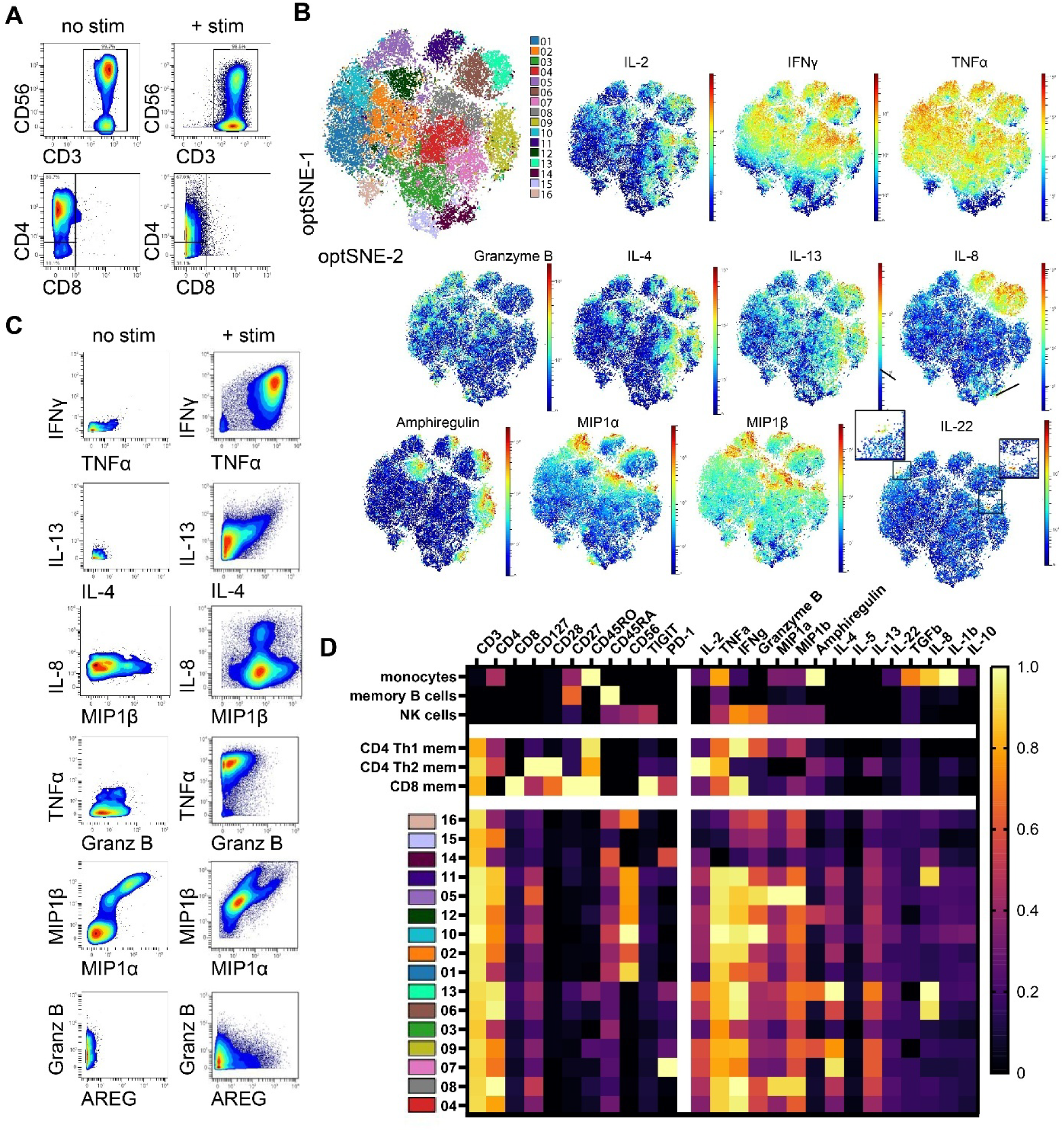
CyTOF analysis of iCD4+ T cells. **(A)** Expression profile of CD3, CD56, CD4 and CD8 in the presence and absence of overnight PMA/ionomycin stimulation. **(B)** opt-SNE projections of stimulated iCD4+ T cells overlaid with color-coded Phenograph clusters 1 through 16, as well as with protein expression heatmaps of intracellular analytes **(C)** Bivariate plots of stimulation-specific upregulation of cytokines and other functional proteins in iCD4+ T cells. **(D)** Heatmap showing column-normalized expression of each marker in Phenograph clusters. The first 6 rows show subsets derived from control PBMCs, while the remaining 16 rows correspond to clusters outlined in fig. 3B.

Secretion of cytokines in response to stimulation is key for CD4+ T cell function. Thus, we measured the secretion of cytokines into supernatant after six hours of PMA and ionomycin stimulation from iCD4+ T cells that have undergone a single round of expansion stimulation (day 68) and two or more rounds of expansion (day 140+) (Fig. 4). In line with CyTOF findings, the iCD4+ T cells secreted a similar diversity of cytokines, with inflammatory cytokines detected at the highest level (Fig. 4). Importantly iCD4+ T cells at baseline without stimulation showed minimal cytokine production and responded upon stimulation with robust cytokine release (Fig. 4).

**Fig. 4.**
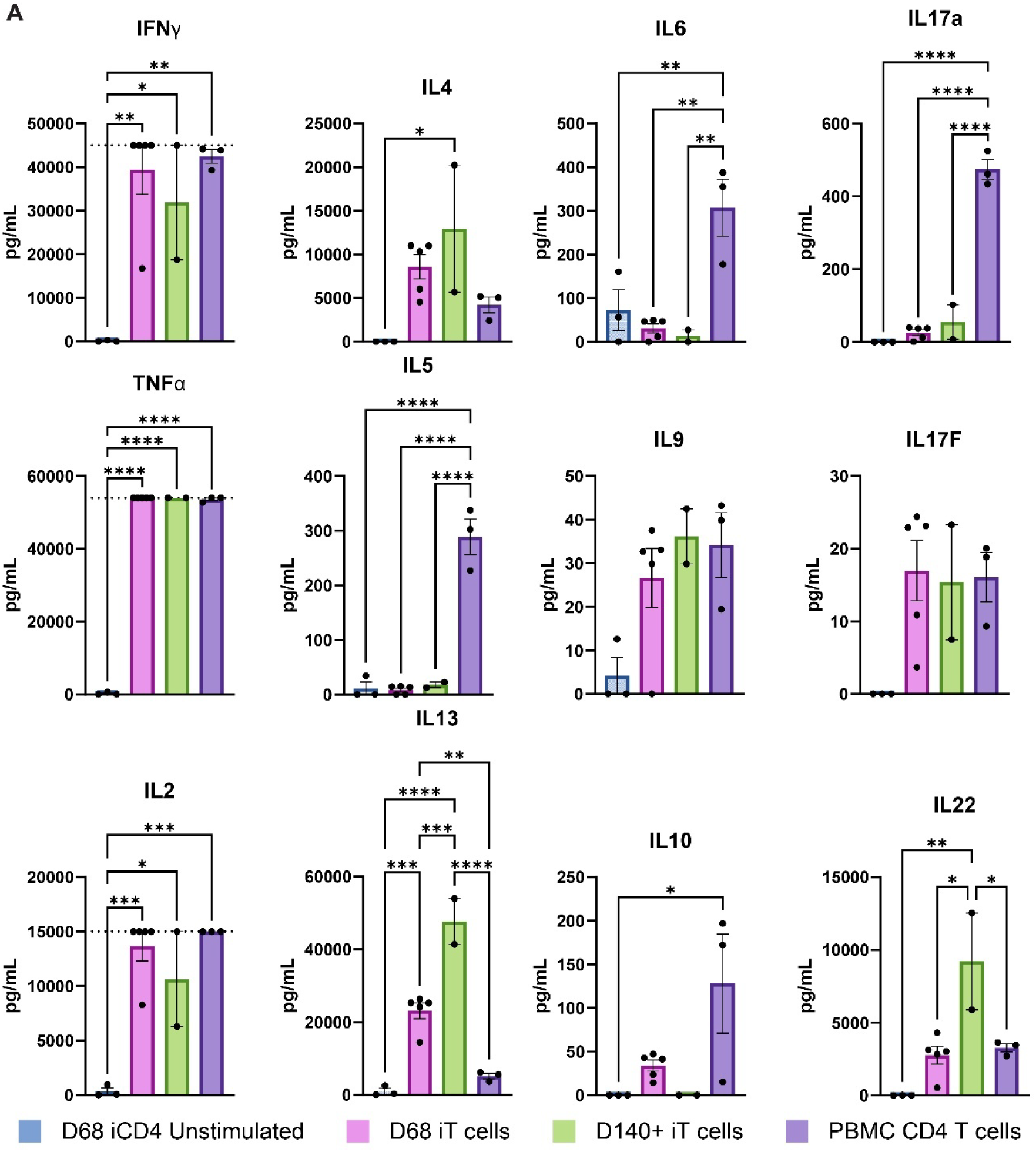
Analysis of cytokine secretion by iCD4+ T cells. (**A**) Concentration of cytokines (pg/ml) in the supernatant of media after six hours of PMA/Ionomycin stimulation. iCD4+ T cells and CD4+ T cells purified from PBMC by magnetic selection were plated in X-Vivo 15 media at 1*10^6 cells/ml. The unstimulated group was rested, while all other groups were treated with PMA (25 ng/ml) and ionomycin (1 µg/ml) at 37 degrees C for 6 hours. Supernatant was collected and analyzed in technical duplicate by 12-Plex LEGENDplex™ HU Th Cytokine Panel (Biolegend) on a Cytek Aurora. Concentration of cytokines were determined using a standard curve by LEGENDplex software (Biolegend). Dotted line indicates the upper limit of detection for the cytokine. N = 3, 5, 2, and 3 differentiations for unstimulated D68 iT cells, D68 iT cells, D140+ iT cells and PBMC T cells respectively. Data shown are group means +/- SEM analyzed by one-way ANOVA with Tukey’s multiple comparison test, *P < 0.05; **P < 0.01; ***P < 0.001; ****P < 0.0001.

### Single-cell RNA sequencing demonstrates the CD4+ helper identity of iCD4+ T cells

To further demonstrate that the iCD4+ T cells show similarity to conventional TCRab T cells, we performed scRNAseq. We chose to sequence iT cells at day 71 and day 103 of culture, alongside CD3+ cells isolated from PBMC. Primary T cells were rested overnight in conditions identical to the iT cells and then prepared for sequencing (Fig. 5A). After sequencing, we performed shared nearest neighbor clustering at 0.15 resolution (Fig. 5B).

**Fig. 5.**
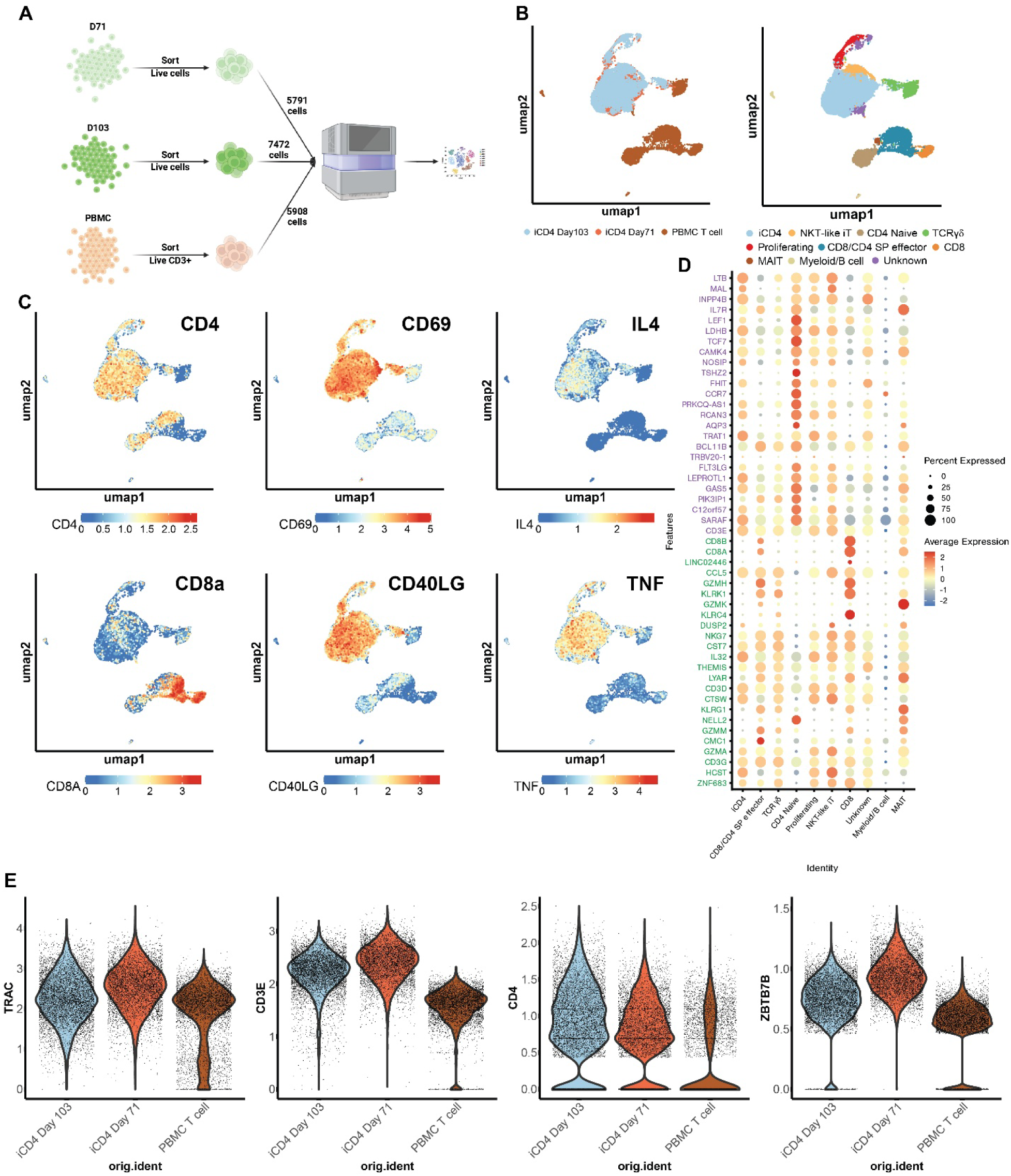
scRNA sequencing reveals characteristic CD4 transcripts in iCD4+ T cells. **(A)** Schematic of scRNAseq experiment. Live cells were purified from iCD4+ T cells cultures at day 71 and 103 of culture by magnetic depletion of debris and dead cells. CD3+ T cells were isolated from PBMC by magnetic selection. **(B)** UMAP projections overlaid with color-coded sample identity (left) derived from Louvain clustering at resolution 0.15 where each color represents annotated clusters. Characteristic marker genes used for annotation indicated in fig S4C. **(C)** UMAP colored by gene expression for key T cell identity, activation and functional genes. **(D)** Bubble plot displaying expression of top 25 DEGs for CD4+ (violet) and CD8+ (green) T cells from literature in each cluster previously defined in B, genes listed in order of highest to lowest significance from the source DEG analysis. (**E**) Key T cell and CD4+ specific marker and transcription factor gene expression data was imputed using ALRA and displayed using a violin plot for each original identity.

UMAP projection followed by Louvain clustering revealed that despite the iCD4+ T cells being cultured for an additional month with an additional round of stimulation, there was almost complete overlap between day 71 and 103 cells. Not surprisingly, the PBMC and iT cell samples clustered separately (Fig. 5B and S6A). Analysis of the differentially expressed genes between the PBMC and iT samples showed that many key drivers behind the differences corresponded to activation states, including CD69 expression and a loss of CD62L which mirrored the flow cytometry results for those markers at day 68 (Fig 5C and S6B) ^47^. There was also significant upregulation of CD40L, classically considered to be a marker of CD4+ T cell activation and differentiation (Fig. 5C)^48,49^.

To further classify the identity of our cells, we used DEG lists generated from a large-scale sequencing study of human PBMCs ^50^. By displaying the top 25 genes from the CD4+ and CD8+ T cell subsets, we observed more similarity to conventional CD4 single positive TCRαβ as compared to conventional CD8 single positive cells (Fig. 5D, S7)^50^. This can also be reflected by clusters three and six which correspond to CD4 and CD8 single positive T cells respectively. Using more specific gene lists and expression of key transcription factors we can identify a CD4+ naïve population in the PBMC sample as well as what appears to be contaminating B and macrophages, MAIT cells, CD8 cells and a mixed CD4/CD8 SP cluster with characteristics of effector cells (Fig. S6C)^50–52^. One mixed cluster of iT and PBMC cells likely corresponded to TCRγδ cells as they showed expression of the TCRδ constant component (Fig. S6C)^50^.

Finally, iCD4+ T cells showed strong expression of *ZBTB7B* (ThPOK), *CD3, TRAC,* and, most importantly, *CD4* with minimal *CD8A* and *CD8B* in the iT samples (Fig. 5E). We also observed the expression of *TNF* and *IL4* across many of the iT cells, further highlighting the functional potential of the cells even without PMA/ionomycin stimulation conditions^47^.

## DISCUSSION

We herein describe the robust generation of functional CD4 single positive T cells from human iPSC in a feeder-free manner accomplished via the removal of Notch signal during TCR mediated DP to SP transition. The resulting iCD4+ T cells proliferate and produce cytokines in response to stimulation. While it is known that a combination of strength and duration of T cell receptor signal influences T lineage divergence, the downstream effects of Notch ligand withdrawal are currently still in question, particularly regarding iPSC derived T cells grown in the presence of high concentrations of Notch ligands^42,53–56^. We found that the absence of Notch ligands allowed for access to the CD4 lineage, an observation that hints at possible mechanisms which have been blocking access to the CD4 lineages in iPSC T cells. Indeed, this also provides an explanation why until now it has not been possible to access the CD4 lineage from iPSCs, as most other protocols rely on T cell maturation conditions containing Notch signaling. We theorize that strong Notch signaling is potentially acting to modulate the strength of signal through the T cell receptor in this system, a theory that warrants future mechanistic studies^57,58^.

CD62L/CCR7 double positivity in a fraction of iCD4+ T cells at day 54 of culture indicates the likelihood of naive T cells present in our iCD4+ T cell cultures at this timepoint. After emigrating from the thymus T cells are known to continue to develop, spending up to 3 weeks maturing further from a recent thymic emigrant to a naïve state. After stimulation through the TCR, naive CD4+ T cells differentiate into memory cells with distinct functional lineages^59,60^. Our next steps will include delineating when within our iPSC to iCD4+ T cell platform naive or naive-like cells can be observed. The use of culture conditions containing alternate cytokines, such as IL-7, IL-15 or both in post-specification culture while removing IL-2, may provide ideal conditions for the maintenance or even expansion of any naive cells present in our cultures. The scRNAseq indicates that after long-term culture with expansion periods, iCD4+ T cells are likely becoming more similar to an effector phenotype as reflected by upregulation of CD69, CD40 ligand and downregulation of CCR7. Eventually, treatment of naive cells with various cytokines may polarize iCD4+ T cells into specific helper subsets, serving as an in vitro screening platform to reveal the molecular pathways responsible for their specification. In this regard, T regulatory cells can be induced from naive T cells through treatment with TGF-β, IL-2, retinoic acid and rapamycin during CD3/CD28 and can act in a highly anti-inflammatory manner^61^.

Alternative methods of generating CD4+ helper T cells from iPSCs have recently been described^62,63^. One used an artificial thymic organoid (ATO) model which consists of a mix of Notch ligand expressing feeder cells as well as iPSC derived lymphoid progenitor cells to create an organoid which is capable of producing CD4+ iPSC T cells^63,64^. The second used a feeder-free system for the generation of CD4+ iPSC T cells using low concentration PMA and ionomycin^62^. The mechanism behind the generation of CD4 SP T cells in the ATO remains unidentified, however it is possible that modulation of Notch signal in the context of the ATO model is different to CD4+ T cell generation in a feeder-free context. The ATO model provides important insights by allowing developing DP cells to have high numbers of cell-cell contacts throughout the selection process. Our platform, in contrast, focuses on CD4+ T cell generation in a feeder-free context solving a critical issue of important practical implications, which is its capability for scalability and control of the final cell product. The second method is feeder-free but uses chemical stimulation of the developing T cells and does not generate CD4 cells expressing TNF and IFNγ. We see clear inflammatory phenotypes including interferon γ within cells generated by our protocol, serving as a potential base for future immunotherapy approaches. Further comparison of each of these methods focusing on deep characterization of the CD4+ iPSC T cells is warranted.

In sum, here we report a scalable, efficient method for the generation of iPSC-derived CD4+ T cells with deep characterization of functional and expression profiles showing similarity to primary CD4+ T cells. iCD4+ T cells may open the gateway for the advancement of iPSC-derived T cell therapies to treat a variety of diseases, particularly those fueled by chronic inflammation.

## METHODS

### Tissue Culture conditions and cell lines

Human iPSC lines were maintained on 6-well tissue culture treated plates coated with hESC-qualified Matrigel (Corning) in 2mL of mTeSR+ (Stemcell technologies) media with added Primocin. Media was changed daily, excepting weekends, where cells were fed twice the standard amount of mTeSR+. Cells were passaged weekly using ReLeSR (Stemcell Technologies) according to manufacturer protocols at a ratio of 1:20 to 1:200. iPSCs were frozen using mFreSR (Stemcell Technologies) and thawed using a ThawSTAR® CFT2 Automated Thawing System. Cells were grown at 37°C, 5% CO2 in standard incubators. iPSC lines used in this study were BU1c2 (XY, EF1a-hSTEMCCA4 loxp lentiviral infection, Cre-excised), BU1c2 TetOn:NICD1, BU2-15-Cr10 TetOn:NICD1 (XY, EF1a-hSTEMCCA4 loxp lentiviral infection, Cre-excised), and BU7 (XX, Sendai virus reprogrammed). All iPSC lines have no known disease-causing mutations and were karyotyped after generation or insertion of the NIDC1 construct by WiCell. iPSC lines are maintained as part of a central bank at Boston University Center for Regenerative Medicine and further information on the derivation and characterization including a karyotype reports can be found at stemcellbank.bu.edu. iPSC lines were thawed between passage 25 and 38 and all differentiations were performed between the second and tenth passage from thaw. iPSC cultures were disposed of if any changes in growth rates were observed. Generation of TetOn:NICD1 lines outlined in prior work ^22^. Human PBMC cells were isolated by density gradient centrifugation with Ficoll and maintained in X-Vivo 15 with 50 IU IL-2 added in T75 or larger flasks.

All cells used in the course of this study were maintained using media containing Primocin (Invivogen) for bacteriostasis and fungistasis. Quarantine and testing of new cell lines in a separate cell culture space to prevent mycoplasma contamination were observed, with three times per year testing of all cells in culture to confirm absence of mycoplasma.

### Differentiation from iPSC to double positive T cells

iPSC cells are differentiated to hematopoietic stem and progenitor cells as previously described over a 12-day period ^22^. CD34+ progenitors are isolated from all floating cells by MACS separation using Miltenyi CD34+ selection kit and LS columns (Miltenyi). CD34+ progenitors are seeded at 5*10^4^ onto TC untreated plates coated with Stemspan Lymphoid Differentiation Coating Material as per manufacturer instructions (StemCell Technologies). The progenitors are fed with Stemspan SFEM II media (StemCell Technologies) supplemented with Lymphoid Progenitor Expansion Supplement (StemCell Technologies) and Primocin. After initial seeding in 1 mL of expansion media cells are cultured for 3 days (Day 15), then fed with an additional 1 mL of expansion media. At Day 19 half of the media is aspirated from each well and replaced with fresh media, then cells are moved to a plate with new coating. Half media changes are performed until day 26, when cells are harvested and re-seeded at 0.5 to 1 * 10^6 cells per mL on new plates coated with Stemspan Lymphoid Differentiation Coating Material in Stemspan SFEM II media supplemented with Lymphoid Progenitor Maturation Supplement (StemCell Technologies) and Primocin. After 3 days maturation media was added as with expansion culture. Half media changes are then performed with maturation media every 3 or 4 days until day 40, when double positive cells were harvested.

### Differentiation from DP T cells to CD4 SP T cells and SP T cell culture conditions

DP cells were plated on 12 well untreated plates coated with 10 ug of RetroNectin (Takara Bio) at variable density from 1*10^5 to 1*10^6 cells per mL in Stemspan SFEM II media supplemented with Lymphoid Progenitor Maturation Supplement (StemCell Technologies) and Primocin. At the time of plating media is supplemented with an additional 10 ng/mL of IL7 and 5 ug/mL of purified anti-CD3 antibody clone UCHT1 or clone OKT3. The media is then changed as outlined previously with half media changes of Stemspan SFEM II media supplemented with Lymphoid Progenitor Maturation Supplement (StemCell Technologies), 10 ng/mL of IL7 and Primocin. At day 54 (14 days after initiation of DP to SP transition) SP T cells can be harvested.

SP iCD4 T cells are moved to a freshly coated RetroNectin plate, in Stemspan SFEM II media supplemented with Lymphoid Progenitor Maturation Supplement (StemCell Technologies), 200 U IL-2 and primocin. Expansion is initiated with the addition of 25 µg/mL of anti CD3/CD28 Immunocult human T cell activator (StemCell Technologies). Through the expansion process cells are fed with half media changes as previously described.

At day 68 iT cells begin to be adapted to X-Vivo 15 media (Lonza) supplemented with 200 U IL-2 by performing half media changes with X-Vivo 15 media. After this time media is changed every 3 or 4 days by performing a half media change with X-Vivo 15 media supplemented with 200 U IL-2.

iPSC T cells were stained for developmental and functional markers using antibodies provided in table S1 and table S3.

### Conventional Flow cytometry

Single cell suspensions were harvested from cultures and resuspended in PBS. Where a live/dead dye was used the cells are incubated with fixable viability stain 780 (BD Biosciences) at 1:1000 for 30 minutes at room temperature in the dark. Cells were then washed with FACS buffer (2% BSA in PBS) and a master mix consisting of antibodies detailed in Table S3. Samples were incubated for 30 minutes at room temperature in the dark. All antibodies were used at 1:100 concentration. UltraComp eBeads (Thermo Fisher) were used for single color controls. Stained samples were read on a Stratedigm instrument equipped with 405, 488, 552 and 640nm lasers.

Analysis performed in FlowJo v10.8 software (BD Biosciences).

### Spectral flow cytometry

Samples were prepared and stained as outlined in flow cytometry. Master mix contained 1x human Fc block (BD Biosciences) and 1x Brilliant Stain Buffer (BD Biosciences) following manufacturer instructions. All antibodies were titrated and used at optimal concentration to minimize background staining while retaining signal. After staining samples were fixed using fixation buffer (BioLegend), washed twice with FACS buffer and resuspended in FACS buffer for analysis. Samples were read on a 5-laser Cytek Aurora instrument and unmixed with SpectraFlo (Cytek) software ordinary least square algorithm. Data analysis was performed in FlowJo v10.8 software (BD Biosciences).

### CyTOF

iPSC derived CD4+ cells were harvested on day 68 and a healthy cryopreserved human PBMC control was thawed. Cells were resuspended in cRPMI at a concentration of 1x10^6^ cells/mL and plated into a 24W Cell-Repellent Surface Plate at 1 mL per well. Cells were rested for 2 hours at 37 °C. Following resting, cells were stimulated with phorbol 12-myristate 13-acetate (PMA) + ionomycin (BioLegend Cell Activation Cocktail, 1:1000 dilution) or left untreated and incubated at 37 °C. After 1 hour, Brefeldin A and monensin (BioLegend, 1:1000 dilution) were added to each well and samples were incubated at 37 °C for an additional 18 hours. Following stimulation, cells were harvested and pooled by sample, counted, and washed with PBS. Up to 3x10^6^ cells per sample were washed with Cell Staining Buffer (Standard BioTools) and incubated with FC-receptor blocking solution (BioLegend). Cells were then incubated with the surface antibody cocktail (see Table S2) in the presence of Cell-ID 103Rh Intercalator (Standard BioTools) for viability staining. After washing with CSB, cells were fixed with Maxpar FIX-I Buffer (Standard BioTools), permeabilized with Maxpar Perm-S Buffer (Standard BioTools) and blocked with sodium heparin blocking solution (Sigma) in Maxpar Perm-S Buffer. Cells were then incubated with the intracellular antibody cocktail, washed with CSB and fixed with 1.6% formaldehyde solution in PBS. Cells were resuspended in Maxpar Fix and Perm buffer supplemented with Cell-ID Intercalator Ir (Standard BioTools) and stored at 2–8 °C overnight prior to acquisition. CyTOF samples were acquired on a CyTOF XT system and samples were normalized with EQ™ Six Element Calibration Beads (Standard BioTools) using CyTOF software. Data analysis was conducted using OMIQ (Dotmatics). Briefly, live singlet cell data from a control PBMC and iCD4 samples were asinh transformed and clustered with Phenograph algorithm and projected into opt-SNE space (perplexity = 30, theta = 0.5, opt-SNE endpoint = 5000; PCA pre-initialization embedding). Clusters were color-coded and overlaid on the opt-SNE projection graphs. Each marker’s mean signal intensities within clustered datasets were organized into hierarchically clustered heatmaps.

### TCR beta sequencing

TCR beta sequencing was accomplished using the bulk-immunoprofiling service through Azenta Life Sciences. Azenta Life Sciences uses a bulk-RNA sequencing platform with amplification of the VDJ regions of the beta TCR chains. Snap-frozen cell pellet samples were submitted from day 68 iPSC-derived T cells. Azenta Life Sciences performed RNA extraction, DNase treatment, cDNA reactions, amplification of the TCR chains, and next generation sequencing. Azenta Life Sciences supplied an analysis report showing the TCR diversity and V-J usage of the submitted sample.

### Cytokine quantification

T cell samples were seeded at 1 million cells per mL in X-Vivo 15 media (Lonza) which had not been supplemented with any cytokines. Phorbol 12-myristate 13-acetate and ionomycin were added at 25 ng/mL and 1 µg/mL concentrations respectively. After a 6 hour incubation each well was centrifuged, and supernatants were harvested and stored at -80 degrees Celsius.

Multiplexed analysis of cytokines was performed using a LEGENDplex™ HU Th Cytokine Panel (12-plex) w/ FP V02 kit from Biolegend. Supernatants were thawed on ice, then diluted in 2 parts assay buffer for each part supernatant. Manufacturer instructions were followed to prepare samples, which were then read on a 6-laser Cytek Aurora. Data was then analyzed in LEGENDplex software (Biolegend).

### Statistics

Statistical tests were performed where indicated in the methods and figure captions. Through this study each N refers to a value from unique differentiation, while in the multiplexed analysis of cytokine release 2 technical replicates were analyzed with the mean value constituting 1 N for the final significance calculation. One and Two way ANOVA tests were performed where indicated using Graphpad Prism version 10.3.0 with Tukey’s multiple comparison test, where p < 0.05 was considered significant. Histogram visualizations include group means +/- SEM with individual measures plotted as separate points unless otherwise indicated in the figure caption. Figure 3 CyTOF data heatmap was generated in R Studio and displays column-normalized expression of each marker.

Statistical analysis of RNA sequencing data is performed in R Studio using methods outlined in the single cell RNA sequencing section of Methods.

### Single cell RNA sequencing

T cell samples were prepared at day 71 and 103 of culture by using Miltenyi Biotec’s dead cell removal kit to increase viability. T cells were isolated from peripheral blood mononuclear cells using a CD3 magnetic separation kit (Miltenyi). After magnetic separation all samples had a viability above 90%.

Samples were processed for sequencing by the Boston University Single Cell Sequencing core, using a 10X Genomics 3’v4 kit. Prepared libraries were sequenced using Illumina NextSeq 2000 Next Generation Sequencing, with a P3 flow cell for 100 cycles.

Sequencing files were mapped to the human genome reference (GRCh37) using CellRanger v3.0.2. Seurat v3.2.3 was used for downstream analysis and quality control ^65^. After inspection of the quality control metrics, cells with 15% to 35% of mitochondrial content and <800 detected genes were excluded for downstream analyses. In addition, doublets were also excluded for downstream analysis. We normalized and scaled the unique molecular identifier (UMI) counts using the regularized negative binomial regression (SCTransform)^66^. Following the standard procedure in Seurat’s pipeline, we performed linear dimensionality reduction (principal component analysis) and used the top 20 principal components to compute the unsupervised Uniform Manifold Approximation and Projection (UMAP)^67^. For clustering of the cells, we used Louvain algorithm which were computed at a range of resolutions from 1.5 to 0.05 (more to fewer clusters) ^68^. Populations were annotated using Louvain Clustering at a resolution of 0.05. Cell cycle scores and classifications were done using the Seurat’s cell-cycle scoring and regression method ^69^. Cluster specific genes were calculated using MAST framework in Seurat wrapper ^70^. ALRA’s algorithm was used to impute the data, to correct for false zeros counts ^71^.

An online Shiny app has been established to allow interactive, user-friendly visualizations of gene expression in each population, https://crem-bu.shinyapps.io/24_09_03_Julian/ along with https://crem-bu.shinyapps.io/24_09_03_Julian_imputed/ for the imputed data^72^.

## ACKNOWLEDGEMENTS

We thank the Boston University Chobanian & Avedisian School of Medicine (BUSM) flow cytometry core and the Boston University Microarray and Sequencing Core for feedback and assistance in the course of this study. We thank the Rhode Island Hospital Stem Cells and Aging (SCA) COBRE Flow Cytometry Core for assistance with collecting CyTOF data.

## Author contributions

Conceptualization: JA, GM

Methodology: JA, DH, MY, ACB, JC, GM Investigation: JA, MY, CMK, GMS, LP Visualization: JA, PSB, FW, ACB Validation: JA, CMK

Funding acquisition: GM Supervision: GM

Writing – original draft: JA, GM

Writing – review & editing: MWY, CMK, PSB, ACB, JC, GM

## Data and materials availability

Single cell RNA sequencing data is available on the GEO repository under accession number GSE279734. All other raw data and code can be provided upon request.

## REFERENCES

1. Joachims, M.L., Chain, J.L., Hooker, S.W., Knott-Craig, C.J., and Thompson, L.F. (2006). Human αβ and γδ Thymocyte Development: TCR Gene Rearrangements, Intracellular TCRβ Expression, and γδ Developmental Potential—Differences between Men and Mice. The Journal of Immunology 176, 1543–1552. 10.4049/jimmunol.176.3.1543.

2. Weerkamp, F., Pike-Overzet, K., and Staal, F.J.T. (2006). T-sing progenitors to commit. Trends in Immunology 27, 125–131. 10.1016/j.it.2006.01.006.

3. Hong, C., Luckey, M.A., and Park, J.-H. (2012). Intrathymic IL-7: The where, when, and why of IL-7 signaling during T cell development. Seminars in Immunology 24, 151–158. 10.1016/j.smim.2012.02.002.

4. Anderson, G., and Takahama, Y. (2012). Thymic epithelial cells: working class heroes for T cell development and repertoire selection. Trends in Immunology 33, 256–263. 10.1016/j.it.2012.03.005.

5. Turka, L.A., Schatz, D.G., Oettinger, M.A., Chun, J.J.M., Gorka, C., Lee, K., McCormack, W.T., and Thompson, C.B. (1991). Thymocyte Expression of RAG-1 and RAG-2: Termination by T Cell Receptor Cross-Linking. Science 253, 778–781. 10.1126/science.1831564.

6. Kumar, B.V., Connors, T.J., and Farber, D.L. (2018). Human T Cell Development, Localization, and Function throughout Life. Immunity 48, 202–213. 10.1016/j.immuni.2018.01.007.

7. Klein, L., Kyewski, B., Allen, P.M., and Hogquist, K.A. (2014). Positive and negative selection of the T cell repertoire: what thymocytes see (and don’t see). Nature Reviews Immunology 14, 377–391. 10.1038/nri3667.

8. Duke-Cohan, J.S., Akitsu, A., Mallis, R.J., Messier, C.M., Lizotte, P.H., Aster, J.C., Hwang, W., Lang, M.J., and Reinherz, E.L. (2023). Pre-T cell receptor self-MHC sampling restricts thymocyte dedifferentiation. Nature 613, 565–574. 10.1038/s41586-022-05555-7.

9. Ross, J.O., Melichar, H.J., Au-Yeung, B.B., Herzmark, P., Weiss, A., and Robey, E.A. (2014). Distinct phases in the positive selection of CD8+ T cells distinguished by intrathymic migration and T-cell receptor signaling patterns. Proceedings of the National Academy of Sciences 111, E2550–E2558. 10.1073/pnas.1408482111.

10. Irla, M. (2022). Instructive Cues of Thymic T Cell Selection. Annual Review of Immunology 40, 95–119. 10.1146/annurev-immunol-101320-022432.

11. Karimi, M.M., Guo, Y., Cui, X., Pallikonda, H.A., Horková, V., Wang, Y.-F., Gil, S.R., Rodriguez-Esteban, G., Robles-Rebollo, I., Bruno, L., et al. (2021). The order and logic of CD4 versus CD8 lineage choice and differentiation in mouse thymus. Nature Communications 12, 99–99. 10.1038/s41467-020-20306-w.

12. Martinez, R.J., and Hogquist, K.A. (2023). The role of interferon in the thymus. Current Opinion in Immunology 84, 102389–102389. 10.1016/j.coi.2023.102389.

13. Park, J.H., Adoro, S., Guinter, T., Erman, B., Alag, A.S., Catalfamo, M., Kimura, M.Y., Cui, Y., Lucas, P.J., Gress, R.E., et al. (2010). Signaling by intrathymic cytokines, not T cell antigen receptors, specifies CD8 lineage choice and promotes the differentiation of cytotoxic-lineage T cells. Nature Immunology 11, 257–264. 10.1038/ni.1840.

14. Xing, Y., Wang, X., Jameson, S.C., and Hogquist, K.A. (2016). Late stages of T cell maturation in the thymus involve NF-κB and tonic type i interferon signaling. Nature Immunology 17, 565–573. 10.1038/ni.3419.

15. Tani-ichi, S., Shimba, A., Wagatsuma, K., Miyachi, H., Kitano, S., Imai, K., Hara, T., and Ikuta, K. (2013). Interleukin-7 receptor controls development and maturation of late stages of thymocyte subpopulations. Proceedings of the National Academy of Sciences 110, 612–617. doi:10.1073/pnas.1219242110.

16. Lonez, C., and Breman, E. (2024). Allogeneic CAR-T Therapy Technologies: Has the Promise Been Met? Cells 13. 10.3390/cells13020146.

17. Worel, N., Grabmeier-Pfistershammer, K., Kratzer, B., Schlager, M., Tanzmann, A., Rottal, A., Körmöczi, U., Porpaczy, E., Staber, P.B., Skrabs, C., et al. (2023). The frequency of differentiated CD3+CD27-CD28- T cells predicts response to CART cell therapy in diffuse large B-cell lymphoma. Frontiers in Immunology 13. 10.3389/fimmu.2022.1004703.

18. Turtle, C.J., Hanafi, L.A., Berger, C., Gooley, T.A., Cherian, S., Hudecek, M., Sommermeyer, D., Melville, K., Pender, B., Budiarto, T.M., et al. (2016). CD19 CAR-T cells of defined CD4+:CD8+ composition in adult B cell ALL patients. J Clin Invest 126, 2123–2138. 10.1172/jci85309.

19. Galli, E., Bellesi, S., Pansini, I., Di Cesare, G., Iacovelli, C., Malafronte, R., Maiolo, E., Chiusolo, P., Sica, S., Sorà, F., and Hohaus, S. (2023). The CD4/CD8 ratio of infused CD19-CAR-T is a prognostic factor for efficacy and toxicity. British Journal of Haematology 203, 564–570. 10.1111/bjh.19117.

20. Xue, D., Lu, S., Zhang, H., Zhang, L., Dai, Z., Kaufman, D.S., and Zhang, J. (2023). Induced pluripotent stem cell-derived engineered T cells, natural killer cells, macrophages, and dendritic cells in immunotherapy. Trends in Biotechnology 41, 907–922. 10.1016/j.tibtech.2023.02.003.

21. Furukawa, Y., Hamano, Y., Shirane, S., Kinoshita, S., Azusawa, Y., Ando, J., Nakauchi, H., and Ando, M. (2022). Advances in Allogeneic Cancer Cell Therapy and Future Perspectives on “Off-the-Shelf” T Cell Therapy Using iPSC Technology and Gene Editing. Cells 11, 269.

22. Heinze, D., Park, S., McCracken, A., Haratianfar, M., Lindstrom, J., Villacorta-Martin, C., Mithal, A., Wang, F., Yang, M.W., Murphy, G., and Mostoslavsky, G. (2022). Notch activation during early mesoderm induction modulates emergence of the T/NK cell lineage from human iPSCs. Stem Cell Reports 17, 2610–2628. 10.1016/j.stemcr.2022.10.007.

23. Takahashi, K., and Yamanaka, S. (2006). Induction of Pluripotent Stem Cells from Mouse Embryonic and Adult Fibroblast Cultures by Defined Factors. Cell 126, 663–676. 10.1016/j.cell.2006.07.024.

24. Wang, B., Iriguchi, S., Waseda, M., Ueda, N., Ueda, T., Xu, H., Minagawa, A., Ishikawa, A., Yano, H., Ishi, T., et al. (2021). Generation of hypoimmunogenic T cells from genetically engineered allogeneic human induced pluripotent stem cells. Nature Biomedical Engineering 5, 429–440. 10.1038/s41551-021-00730-z.

25. Maeda, T., Nagano, S., Ichise, H., Kataoka, K., Yamada, D., Ogawa, S., Koseki, H., Kitawaki, T., Kadowaki, N., and Takaori-Kondo, A. (2016). Regeneration of CD8αβ T cells from T-cell–derived iPSC imparts potent tumor antigen-specific cytotoxicity. Cancer research 76, 6839–6850.

26. Kruse, B., Buzzai, A.C., Shridhar, N., Braun, A.D., Gellert, S., Knauth, K., Pozniak, J., Peters, J., Dittmann, P., Mengoni, M., et al. (2023). CD4+ T cell-induced inflammatory cell death controls immune-evasive tumours. Nature 618, 1033–1040. 10.1038/s41586-023-06199-x.

27. Boulch, M., Cazaux, M., Loe-Mie, Y., Thibaut, R., Corre, B., Lemaître, F., Grandjean, C.L., Garcia, Z., and Bousso, P. (2021). A cross-talk between CAR T cell subsets and the tumor microenvironment is essential for sustained cytotoxic activity. Sci Immunol 6. 10.1126/sciimmunol.abd4344.

28. Melenhorst, J.J., Chen, G.M., Wang, M., Porter, D.L., Chen, C., Collins, M.A., Gao, P., Bandyopadhyay, S., Sun, H., Zhao, Z., et al. (2022). Decade-long leukaemia remissions with persistence of CD4+ CAR T cells. Nature 602, 503–509. 10.1038/s41586-021-04390-6.

29. Wang, X., Popplewell, L.L., Wagner, J.R., Naranjo, A., Blanchard, M.S., Mott, M.R., Norris, A.P., Wong, C.W., Urak, R.Z., Chang, W.C., et al. (2016). Phase 1 studies of central memory-derived CD19 CAR T-cell therapy following autologous HSCT in patients with B-cell NHL. Blood 127, 2980–2990. 10.1182/blood-2015-12-686725.

30. Wang, D., Aguilar, B., Starr, R., Alizadeh, D., Brito, A., Sarkissian, A., Ostberg, J.R., Forman, S.J., and Brown, C.E. (2018). Glioblastoma-targeted CD4+ CAR T cells mediate superior antitumor activity. JCI Insight 3. 10.1172/jci.insight.99048.

31. Iriguchi, S., Yasui, Y., Kawai, Y., Arima, S., Kunitomo, M., Sato, T., Ueda, T., Minagawa, A., Mishima, Y., Yanagawa, N., et al. (2021). A clinically applicable and scalable method to regenerate T-cells from iPSCs for off-the-shelf T-cell immunotherapy. Nature Communications 12, 430. 10.1038/s41467-020-20658-3.

32. Montel-Hagen, A., and Crooks, G.M. (2019). From pluripotent stem cells to T cells. Experimental Hematology 71, 24–31. 10.1016/j.exphem.2018.12.001.

33. Ashby, K.M., and Hogquist, K.A. (2023). A guide to thymic selection of T cells. Nature Reviews Immunology. 10.1038/s41577-023-00911-8.

34. Mangelinck, A., Dubuisson, A., Becht, E., Dromaint-Catesson, S., Fasquel, M., Provost, N., Walas, D., Darville, H., Galizzi, J.-P., Lefebvre, C., et al. (2024). Characterization of CD4+ and CD8+ T cells responses in the mixed lymphocyte reaction by flow cytometry and single cell RNA sequencing. Frontiers in Immunology Volume 14-2023. 10.3389/fimmu.2023.1320481.

35. Azzam, H.S., Grinberg, A., Lui, K., Shen, H., Shores, E.W., and Love, P.E. (1998). CD5 Expression Is Developmentally Regulated By T Cell Receptor (TCR) Signals and TCR Avidity. Journal of Experimental Medicine 188, 2301–2311. 10.1084/jem.188.12.2301.

36. Azzam, H.S., DeJarnette, J.B., Huang, K., Emmons, R., Park, C.-S., Sommers, C.L., El-Khoury, D., Shores, E.W., and Love, P.E. (2001). Fine Tuning of TCR Signaling by CD5. The Journal of Immunology 166, 5464–5472. 10.4049/jimmunol.166.9.5464.

37. Li, Y., Guaman Tipan, P., Selden, H.J., Srinivasan, J., Hale, L.P., and Ehrlich, L.I.R. (2023). CCR4 and CCR7 differentially regulate thymocyte localization with distinct outcomes for central tolerance. eLife 12, e80443. 10.7554/eLife.80443.

38. Kwan, J., and Killeen, N. (2004). CCR7 Directs the Migration of Thymocytes into the Thymic Medulla1. The Journal of Immunology 172, 3999–4007. 10.4049/jimmunol.172.7.3999.

39. Fink, P.J. (2013). The Biology of Recent Thymic Emigrants. Annual Review of Immunology 31, 31–50. 10.1146/annurev-immunol-032712-100010.

40. Park, J.E., Botting, R.A., Conde, C.D., Popescu, D.M., Lavaert, M., Kunz, D.J., Goh, I., Stephenson, E., Ragazzini, R., Tuck, E., et al. (2020). A cell atlas of human thymic development defines T cell repertoire formation. Science 367. 10.1126/science.aay3224.

41. Snook, J.P., Kim, C., and Williams, M.A. (2018). TCR signal strength controls the differentiation of CD4+ effector and memory T cells. Science Immunology 3, eaas9103. doi:10.1126/sciimmunol.aas9103.

42. Luckey, M.A., Kimura, M.Y., Waickman, A.T., Feigenbaum, L., Singer, A., and Park, J.-H. (2014). The transcription factor ThPOK suppresses Runx3 and imposes CD4+ lineage fate by inducing the SOCS suppressors of cytokine signaling. Nature Immunology 15, 638–645. 10.1038/ni.2917.

43. Levine, Jacob H., Simonds, Erin F., Bendall, Sean C., Davis, Kara L., Amir, E.-ad D., Tadmor, Michelle D., Litvin, O., Fienberg, Harris G., Jager, A., Zunder, Eli R., et al. (2015). Data-Driven Phenotypic Dissection of AML Reveals Progenitor-like Cells that Correlate with Prognosis. Cell 162, 184–197. 10.1016/j.cell.2015.05.047.

44. Belkina, A.C., Ciccolella, C.O., Anno, R., Halpert, R., Spidlen, J., and Snyder-Cappione, J.E. (2019). Automated optimized parameters for T-distributed stochastic neighbor embedding improve visualization and analysis of large datasets. Nature Communications 10, 5415–5415. 10.1038/s41467-019-13055-y.

45. Alizadeh, D., Wong, R.A., Gholamin, S., Maker, M., Aftabizadeh, M., Yang, X., Pecoraro, J.R., Jeppson, J.D., Wang, D., Aguilar, B., et al. (2021). IFNγ Is Critical for CAR T Cell-Mediated Myeloid Activation and Induction of Endogenous Immunity. Cancer Discov 11, 2248–2265. 10.1158/2159-8290.Cd-20-1661.

46. Boulch, M., Cazaux, M., Cuffel, A., Guerin, M.V., Garcia, Z., Alonso, R., Lemaître, F., Beer, A., Corre, B., Menger, L., et al. (2023). Tumor-intrinsic sensitivity to the pro-apoptotic effects of IFN-γ is a major determinant of CD4+ CAR T-cell antitumor activity. Nature Cancer 4, 968–983. 10.1038/s43018-023-00570-7.

47. Reed, J., and Wetzel, S.A. (2018). CD4(+) T Cell Differentiation and Activation. Methods Mol Biol 1803, 335–351. 10.1007/978-1-4939-8549-4_20.

48. Clarke, S.R.M. (2000). The critical role of CD40/CD40L in the CD4-dependent generation of CD8+ T cell immunity. Journal of Leukocyte Biology 67, 607–614. 10.1002/jlb.67.5.607.

49. Elgueta, R., Benson, M.J., De Vries, V.C., Wasiuk, A., Guo, Y., and Noelle, R.J. (2009). Molecular mechanism and function of CD40/CD40L engagement in the immune system. Immunological Reviews 229, 152–172. 10.1111/j.1600-065X.2009.00782.x.

50. Terekhova, M., Swain, A., Bohacova, P., Aladyeva, E., Arthur, L., Laha, A., Mogilenko, D.A., Burdess, S., Sukhov, V., Kleverov, D., et al. (2024). Single-cell atlas of healthy human blood unveils age-related loss of NKG2C+GZMB-CD8+ memory T cells and accumulation of type 2 memory T cells. Immunity 57, 188–192. 10.1016/j.immuni.2023.12.014.

51. Garner, L.C., Klenerman, P., and Provine, N.M. (2018). Insights Into Mucosal-Associated Invariant T Cell Biology From Studies of Invariant Natural Killer T Cells. Frontiers in Immunology 9.

52. Malarkannan, S. (2020). NKG7 makes a better killer. Nature Immunology 21, 1139–1140. 10.1038/s41590-020-0767-5.

53. Singer, A. (2002). New perspectives on a developmental dilemma: the kinetic signaling model and the importance of signal duration for the CD4/CD8 lineage decision. Current Opinion in Immunology 14, 207–215. 10.1016/S0952-7915(02)00323-0.

54. Zeidan, N., Damen, H., Roy, D.-C., and Dave, V.P. (2019). Critical Role for TCR Signal Strength and MHC Specificity in ThPOK-Induced CD4 Helper Lineage Choice. The Journal of Immunology 202, 3211–3225. 10.4049/jimmunol.1801464.

55. Karimi, M.M., Guo, Y., Cui, X., Pallikonda, H.A., Horková, V., Wang, Y.-F., Gil, S.R., Rodriguez-Esteban, G., Robles-Rebollo, I., Bruno, L., et al. (2021). The order and logic of CD4 versus CD8 lineage choice and differentiation in mouse thymus. Nature Communications 12, 99. 10.1038/s41467-020-20306-w.

56. Shinzawa, M., Moseman, E.A., Gossa, S., Mano, Y., Bhattacharya, A., Guinter, T., Alag, A., Chen, X., Cam, M., McGavern, D.B., et al. (2022). Reversal of the T cell immune system reveals the molecular basis for T cell lineage fate determination in the thymus. Nature Immunology 23, 731–742. 10.1038/s41590-022-01187-1.

57. Izon, D.J., Punt, J.A., Xu, L., Karnell, F.G., Allman, D., Myung, P.S., Boerth, N.J., Pui, J.C., Koretzky, G.A., and Pear, W.S. (2001). Notch1 Regulates Maturation of CD4+ and CD8+ Thymocytes by Modulating TCR Signal Strength. Immunity 14, 253–264. 10.1016/S1074-7613(01)00107-8.

58. Vanderbeck, A., and Maillard, I. (2020). Notch signaling at the crossroads of innate and adaptive immunity. Journal of Leukocyte Biology 109, 535–548. 10.1002/jlb.1ri0520-138r.

59. Cunningham, C.A., Helm, E.Y., and Fink, P.J. (2018). Reinterpreting recent thymic emigrant function: defective or adaptive? Curr Opin Immunol 51, 1–6. 10.1016/j.coi.2017.12.006.

60. Berkley, A.M., Hendricks, D.W., Simmons, K.B., and Fink, P.J. (2013). Recent thymic emigrants and mature naive T cells exhibit differential DNA methylation at key cytokine loci. J Immunol 190, 6180–6186. 10.4049/jimmunol.1300181.

61. Schmidt, A., Éliás, S., Joshi, R.N., and Tegnér, J. (2016). In Vitro Differentiation of Human CD4+FOXP3+ Induced Regulatory T Cells (iTregs) from Naive CD4+ T Cells Using a TGFb-containing Protocol. JoVE, e55015. doi:10.3791/55015.

62. Fong, H., Mendel, M., Jascur, J., Najmi, L., Kim, K., Lew, G., Garimalla, S., Schock, S., Hu, J., Villegas, A.G., et al. (2025). A serum- and feeder-free system to generate CD4 and regulatory T cells from human iPSCs. Stem Cells 43. 10.1093/stmcls/sxaf001.

63. Yano, H., Koga, K., Sato, T., Shinohara, T., Iriguchi, S., Matsuda, A., Nakazono, K., Shioiri, M., Miyake, Y., Kassai, Y., et al. (2024). Human iPSC-derived CD4(+) Treg-like cells engineered with chimeric antigen receptors control GvHD in a xenograft model. Cell Stem Cell 31, 795–802.e796. 10.1016/j.stem.2024.05.004.

64. Montel-Hagen, A., Seet, C.S., Li, S., Chick, B., Zhu, Y., Chang, P., Tsai, S., Sun, V., Lopez, S., Chen, H.-C., et al. (2019). Organoid-Induced Differentiation of Conventional T Cells from Human Pluripotent Stem Cells. Cell Stem Cell 24, 376–389.e378. 10.1016/j.stem.2018.12.011.

65. Stuart, T., Butler, A., Hoffman, P., Hafemeister, C., Papalexi, E., Mauck, W.M., 3rd, Hao, Y., Stoeckius, M., Smibert, P., and Satija, R. (2019). Comprehensive Integration of Single-Cell Data. Cell 177, 1888–1902.e1821. 10.1016/j.cell.2019.05.031.

66. Hafemeister, C., and Satija, R. (2019). Normalization and variance stabilization of single-cell RNA-seq data using regularized negative binomial regression. Genome Biology 20, 296. 10.1186/s13059-019-1874-1.

67. McInnes, L., Healy, J., and Melville, J. (2020). UMAP: Uniform Manifold Approximation and Projection for Dimension Reduction. arXiv. 10.48550/arXiv.1802.03426.

68. Blondel, V.D., Guillaume, J.-L., Lambiotte, R., and Lefebvre, E. (2008). Fast unfolding of communities in large networks. Journal of Statistical Mechanics: Theory and Experiment 2008, P10008. 10.1088/1742-5468/2008/10/P10008.

69. Tirosh, I., Izar, B., Prakadan, S.M., Wadsworth, M.H., 2nd, Treacy, D., Trombetta, J.J., Rotem, A., Rodman, C., Lian, C., Murphy, G., et al. (2016). Dissecting the multicellular ecosystem of metastatic melanoma by single-cell RNA-seq. Science 352, 189–196. 10.1126/science.aad0501.

70. Finak, G., McDavid, A., Yajima, M., Deng, J., Gersuk, V., Shalek, A.K., Slichter, C.K., Miller, H.W., McElrath, M.J., Prlic, M., et al. (2015). MAST: a flexible statistical framework for assessing transcriptional changes and characterizing heterogeneity in single-cell RNA sequencing data. Genome Biol 16, 278. 10.1186/s13059-015-0844-5.

71. Linderman, G.C., Zhao, J., Roulis, M., Bielecki, P., Flavell, R.A., Nadler, B., and Kluger, Y. (2022). Zero-preserving imputation of single-cell RNA-seq data. Nature Communications 13,192. 10.1038/s41467-021-27729-z.

72. Ouyang, J.F., Kamaraj, U.S., Cao, E.Y., and Rackham, O.J.L. (2021). ShinyCell: simple and sharable visualization of single-cell gene expression data. Bioinformatics 37, 3374–3376. 10.1093/bioinformatics/btab209.

