## Supplemental figures and tables for "Generation of effector CD4+ T cells from human iPSC"

**Affiliations:**

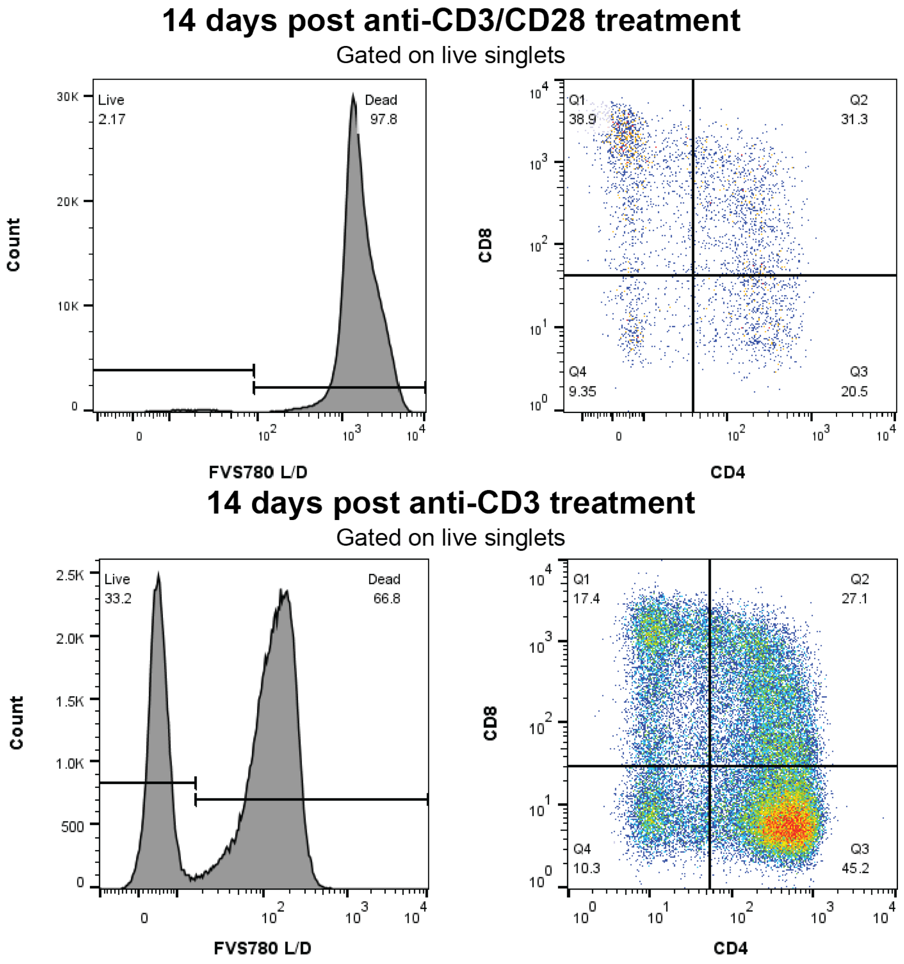

**Fig. S1. Anti-CD3 treatment alone improves viability of iPSC derived CD4+ T cells.** Representative viability and CD4/8 flow cytometry of iT cells generated 14 days after DP iT cells are stimulated with anti-CD3/CD28 (25 ul/ml immunocult, StemCell) or anti-CD3 (5 ug/ml OKT3).

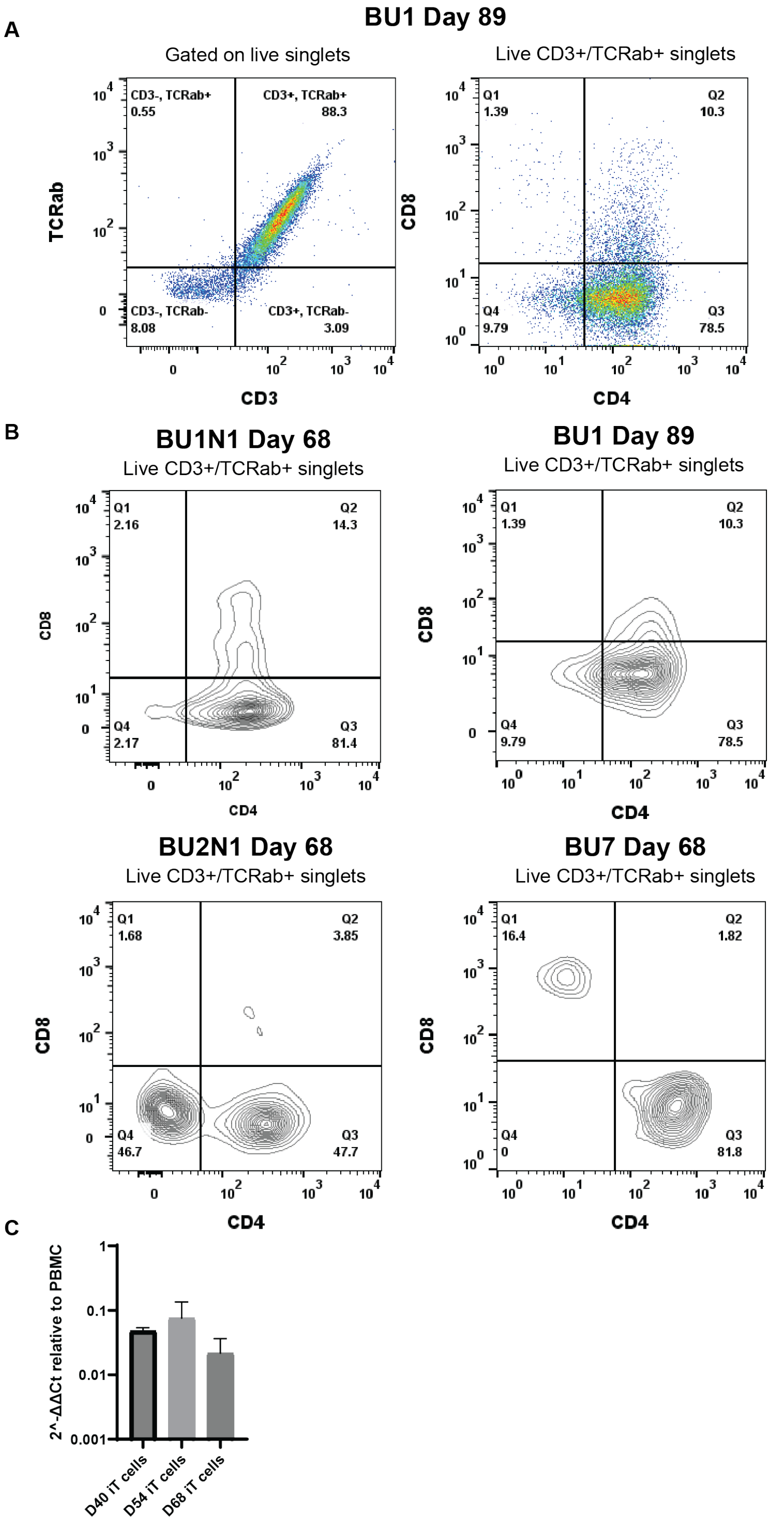

**Fig. S2. CD4 specification in additional iPSC lines with and without inducible NICD1. (A)** Flow cytometry of WT BU1 background iT cells after CD4 specification and 1 round of expansion. **(B)** Flow cytometry of BU1N1, BU1, BU2N1 and BU7 background iT cells after CD4 specification and 1 round of expansion. **(C)** qPCR analysis of ThPOK transcription over 1 differentiation of BU1N1 cells at day 40, 54 and 68, relative to PBMC T cells using ACTb as a housekeeping gene in technical triplicate. Mean + SEM.

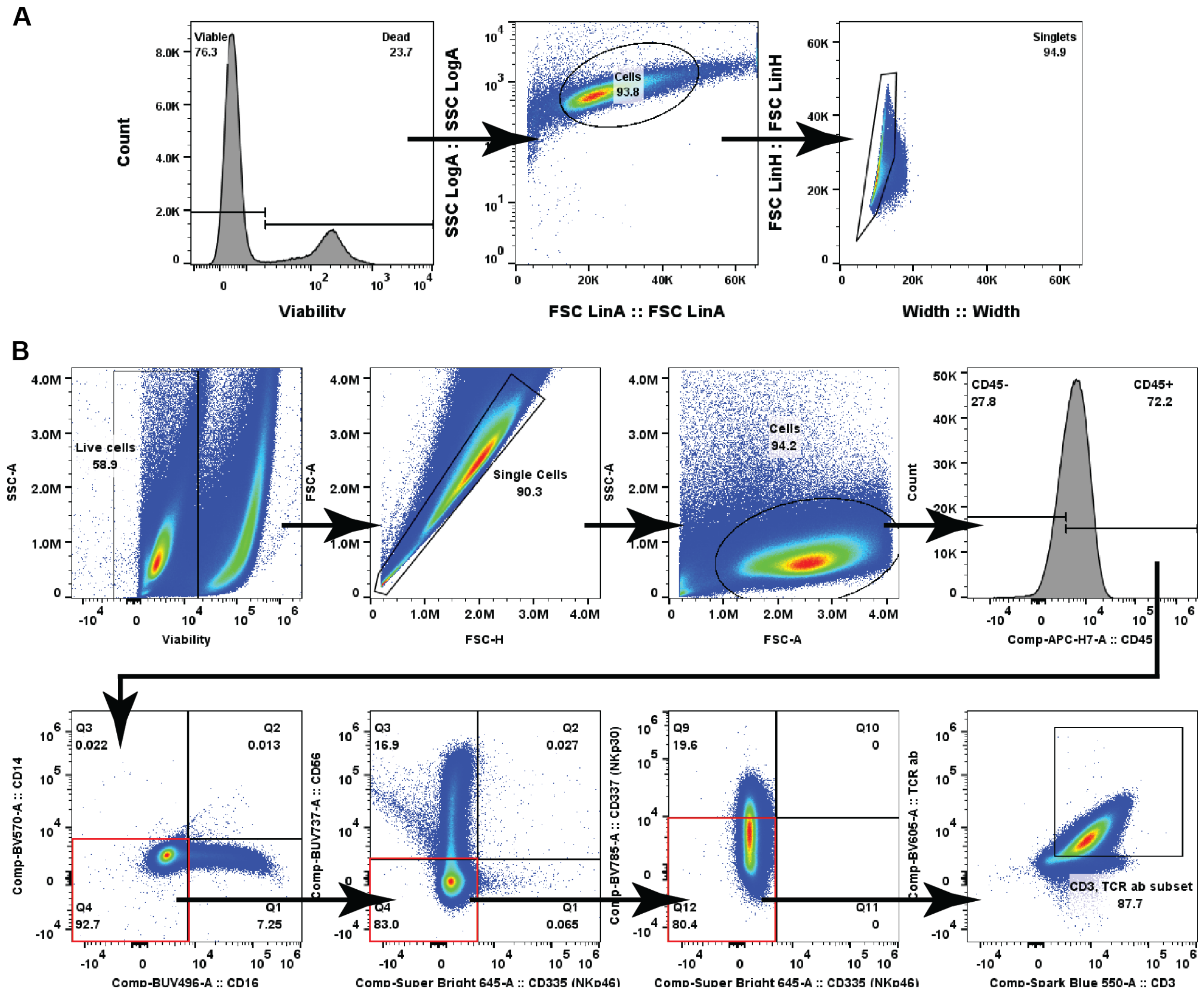

**Fig. S3. Gating strategy for flow cytometry analysis. (A)** Gating strategy for iT cells in figure 1. **(B)** Gating strategy for iT cells in figure 2.

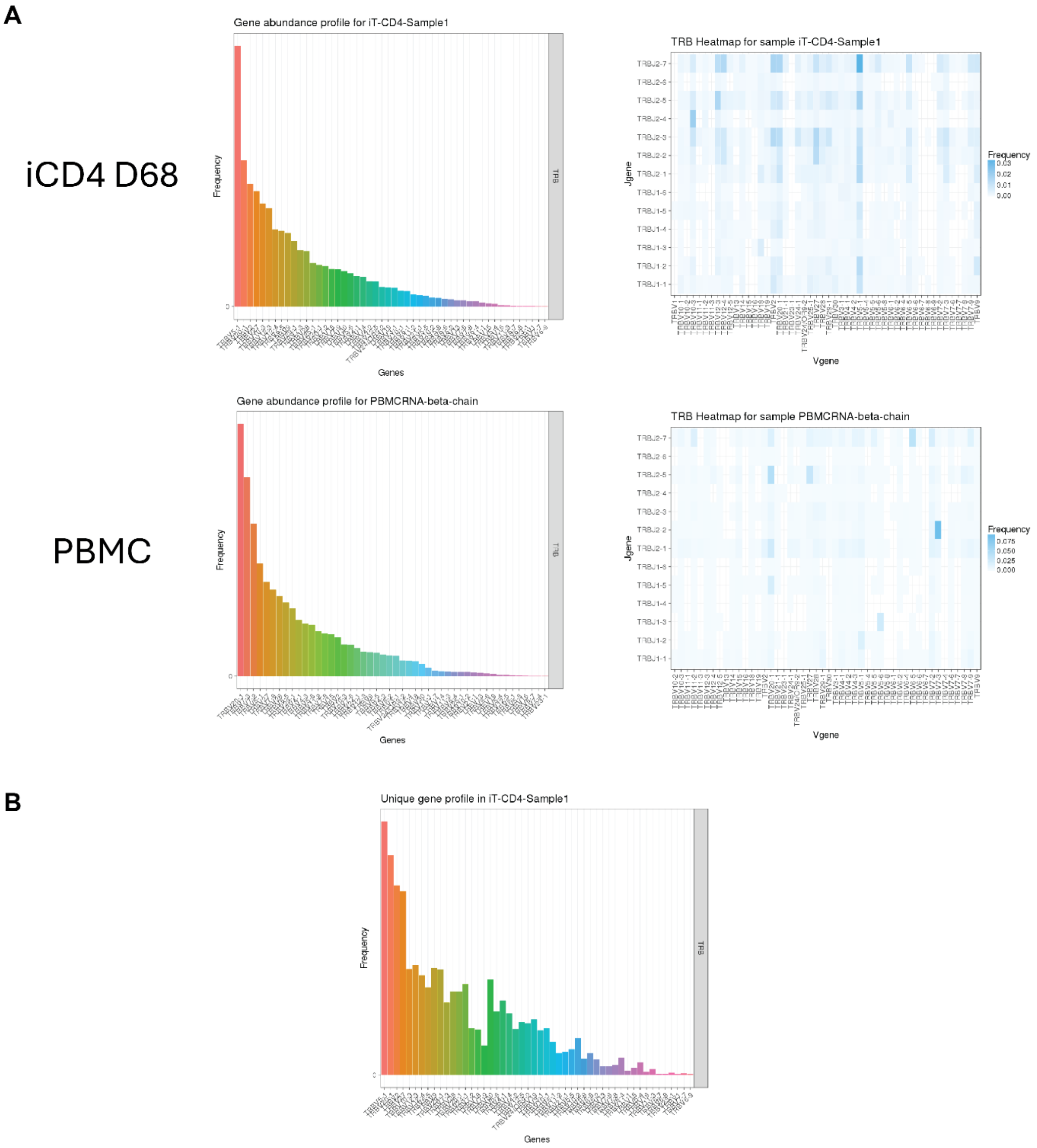

**Fig. S4. TCRb sequencing of iCD4 T cells. (A)** Abundance plots of TRBV genes and heatmaps of TRBV and TRBJ gene usage in both iCD4 cells at day 68 and T cells purified from PBMC. RNA was extracted from snap-frozen cell pellets of one million iCD4 T cells at day 68 of culture and sequenced. **(B)** Unique gene frequency plot of TRBV genes in iCD4 T cell sample.

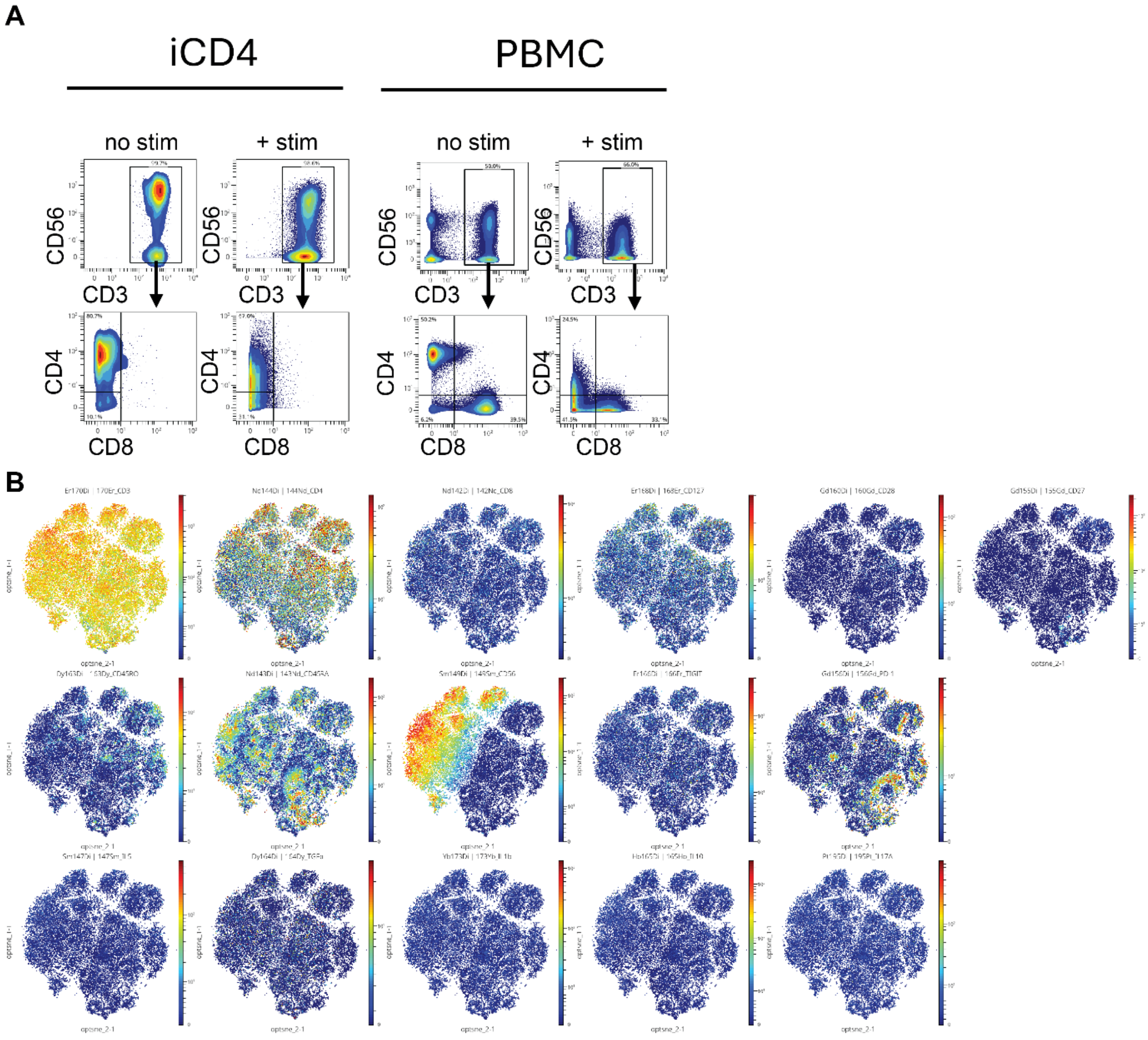

**Fig. S5. Expression of additional proteins by CyTOF (A)** iCD4 comparison to PBMC derived T cells with and without overnight PMA/ionomycin stimulation. **(B)** optSNE map of stimulated iCD4+ T cells colored by expression of additional markers.

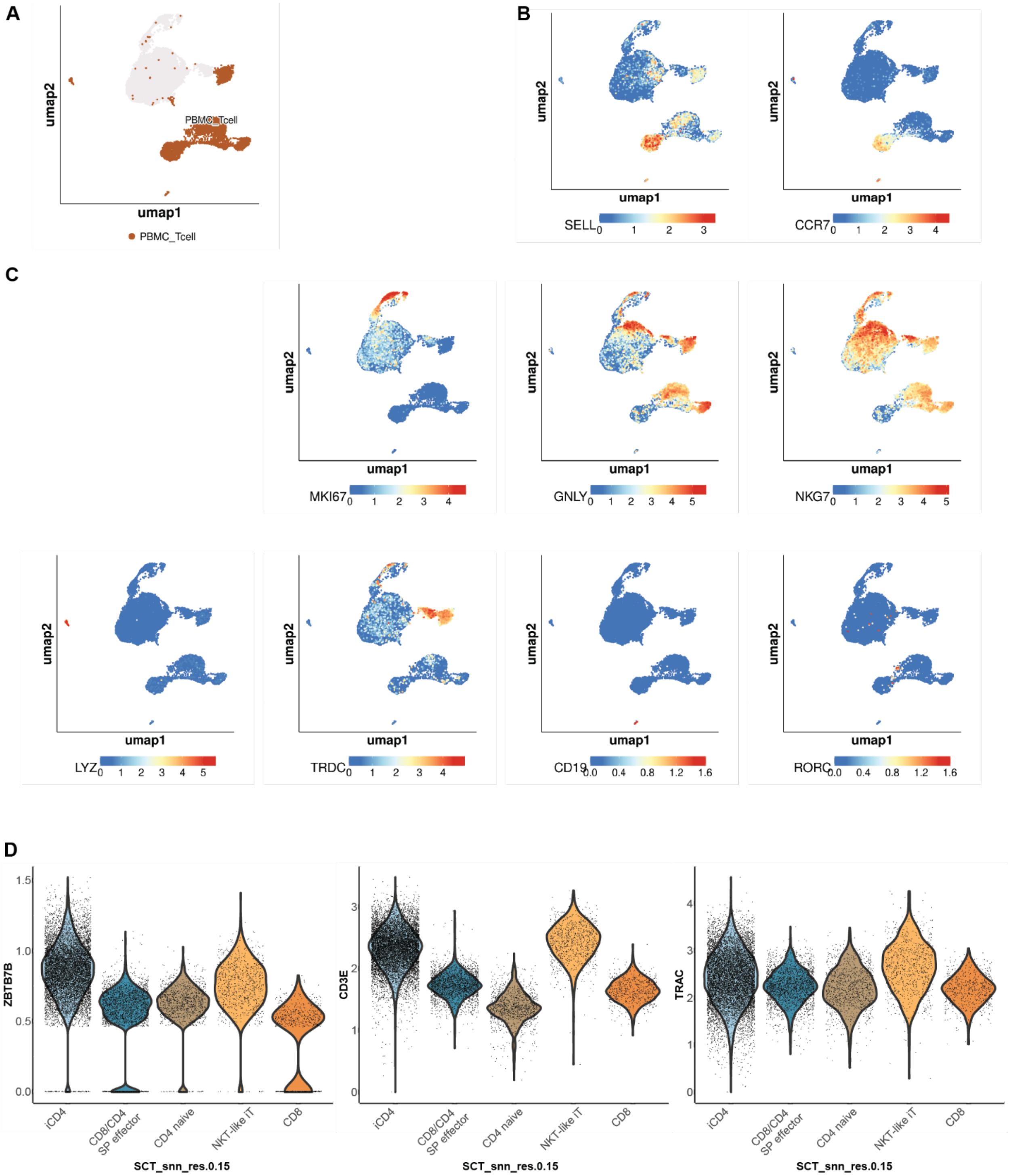

**Fig. S6. Additional genes revealed in scRNAseq for annotation and CD4 identity of iT cells.** **(A)** UMAP showing only PBMC cells. **(B)** UMAP colored by expression of naïve CD4 genes. **(C)** UMAP colored by expression of markers for proliferating cells, NKT cells, myeloid cells, B cells and MAIT cells. **(D)** Violin plot of imputed data for key clusters showing key CD4 T cell gene expression.

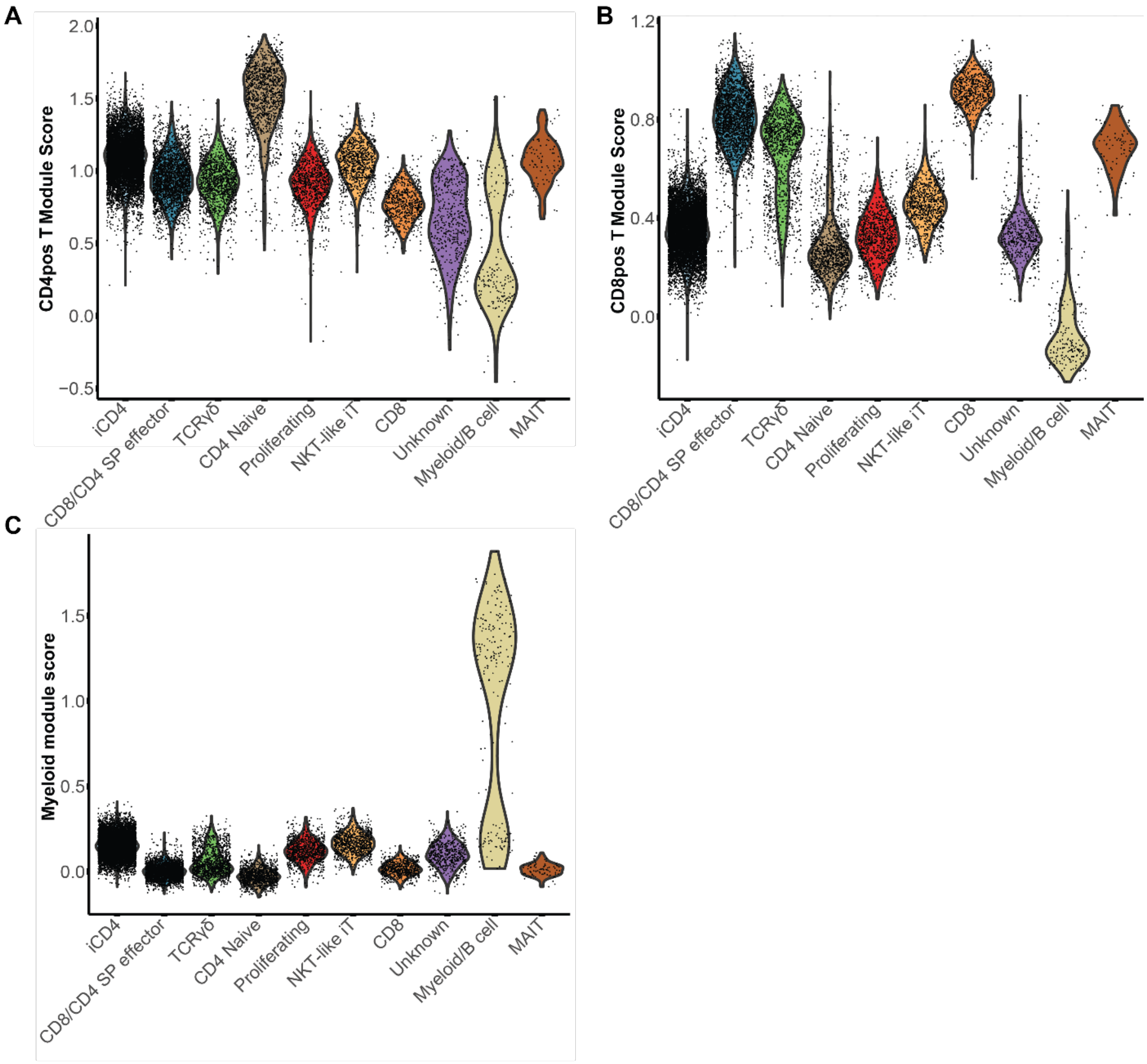

**Fig. S7. Module scores by cluster from scRNAseq experiment.** Module score for CD4+ T cells **(A)**, CD8+ TRAV 1-2- T cells **(B)**, and myeloid cells **(C)**, derived from the top 100 DEGs in these populations from a large scale single cell RNA sequencing study of human PBMC (*50*) plotted by cluster.

| Number | Fluorophore | Antibody | Titer | Catalog number |
|---|---|---|---|---|
| 1 | BUV395 | CD8b | 1:50 | 742396 |
| 2 | BUV496 | CD16 | 1:50 | 612945 |
| 3 | BUV615 | CD62L | 1:400 | 751364 |
| 4 | BUV737 | CD56 | 1:100 | 612767 |
| 5 | BUV805 | CD8a | 1:200 | 612890 |
| 6 | BV421 | CD69 | 1:100 | 310929 |
| 7 | BV480 | CD7 | 1:50 | 566161 |
| 8 | BV510 | CD33 | 1:100 | 366609 |
| 9 | BV570 | CD14 | 1:50 | 301831 |
| 10 | BV605 | TCRab | 1:50 | 306731 |
| 11 | BV711 | CD5 | 1:200 | 563170 |
| 12 | BV785 | NKp30 | 1:50 | 325229 |
| 13 | Spark Blue 550 | CD3 | 1:100 | 344851 |
| 14 | SuperBright 645 | NKp46 CD335 | 1:50 | 64-3351-80 |
| 15 | FITC | TCRgd | 1:50 | 130-114-029 |
| 16 | PE | CD25 | 1:50 | 12-0257-41 |
| 17 | PE-Dazzle594 | MHC-I | 1:800 | 311439 |
| 18 | PE-Cy7 | CCR7 | 1:100 | 567314 |
| 19 | AF647 | CD1a | 1:100 | 300116 |
| 20 | APCH7 | CD45 | 1:800 | 560274 |
| 21 | APC-Fire810 | CD4 | 1:400 | 344661 |
| 22 | PerCP-Cy5.5 | CD24 | 1:25 | 311115 |
| 23 | LD Blue Viability | Viability | 1:1000 | L23105 |

**Table 1.** Panel and titers for spectral flow cytometry.

| Number | Metal | Marker | Clone |
|---|---|---|---|
| 1 | 149 Sm | CD56 | NCAM16.2 |
| 2 | 143 Nd | CD45RA | H100 |
| 3 | 153 Eu | CD25 | BC96 |
| 4 | 155 Gd | CD27 | L128 |
| 5 | 160 Gd | CD28 | CD28.2 |
| 6 | 163 Dy | CD45RO | UCHL1 |
| 7 | 168 Er | CD127 | A019D5 |
| 8 | 164 Dy | TGFb | S20006A |
| 9 | 156 Gd | PD-1 | EH12.2H7 |
| 10 | 166 Er | TIGIT | MBSA43 |
| 11 | | 103Rh Viability | |
| 12 | 110 Cd | IL-3 | BVD3-1F9 |
| 13 | 113 Cd | MIP1a | W16009B |
| 14 | 139 La | IL-8 | G265-8 |
| 15 | 152 Sm | MIP1b | D21-1351 |
| 16 | 162 Dy | Amphiregulin | AREG559 |
| 17 | 173 Yb | IL-1b | CRM56 |
| 18 | 144 Nd | CD4 | SK3 |
| 19 | 112 Cd | IL-2 | MQ1-17H12 |
| 20 | 114 Cd | TNFa | MAb11 |
| 21 | 116 Cd | IFN-y | B27 |
| 22 | 147 Sm | IL-5 | TRFK5 |
| 23 | 150 Nd | IL-22 | HI100 |
| 24 | 165 Ho | IL-10 | JES3-9D7 |
| 25 | 169 Tm | IL-13 | JES10 |
| 26 | 171 Yb | IL-4 | MP4-25D2 |
| 27 | 172 Yb | IL-21 | 3A3-N2 |
| 28 | 195 Pt | IL-17A | BL168 |
| 29 | 198 Pt | Granzyme B | GB11 |
| 30 | 142 Nd | CD8 | SK1 |
| 31 | 170 Er | CD3 | UCHT1 |

**Table 2.** Panel used for CyTOF study.

| **REAGENT or RESOURCE** | **SOURCE** | **IDENTIFIER** |
|---|---|---|
| **Antibodies** | | |
| BV421 Mouse Anti-Human CD34 | BD Bioscience | Cat#562577 |
| PE Mouse Anti-Human CD56 | Biolegend | Cat#304605 |
| BV605 Mouse Anti-Human CD5 | BD Bioscience | Cat#563945 |
| AF647 Mouse Anti-Human HLA-DR, DP, DQ | BD Bioscience | Cat#563591 |
| BV421 Mouse Anti-Human CD1a | BD Bioscience | Cat#563939 |
| BB515 Mouse Anti-Human CD4 | BD Bioscience | Cat#564500 |

| | | |
|---|---|---|
| APC Mouse Anti-Human CD7 | BD Bioscience | Cat#561604 |
| FITC Mouse Anti-Human CD16 | BD Bioscience | Cat#561308 |
| APC Mouse Anti-Human CD69 | BD Bioscience | Cat#560967 |
| BV421 Mouse Anti-Human CD8 | BD Bioscience | Cat#562428 |
| BUV395 Mouse Anti-Human CD8b | BD Biosciences | Cat# 742396 |
| LD Blue Viability | Thermo Fisher | Cat# L23105 |
| BUV496 Mouse Anti-Human CD16 | BD Biosciences | Cat# 612945 |
| BUV737 Mouse Anti-Human CD56 | BD Biosciences | Cat# 612767 |
| BUV805 Mouse Anti-Human CD8 | BD Biosciences | Cat# 612890 |
| BV421 Mouse Anti-Human CD69 | Biolegend | Cat# 310929 |
| FITC Mouse Anti-Human TCR γδ | BD Biosciences | Cat# 559878 |
| BV480 Mouse Anti-Human CD7 | BD Biosciences | Cat# 566161 |
| BV510 Mouse Anti-Human CD33 | Biolegend | Cat# 366609 |
| BV570 Mouse Anti-Human CD14 | Biolegend | Cat# 301831 |
| BV605 Mouse Anti-Human TCR a/b | Biolegend | Cat# 306731 |
| SuperBright 645 Rat Anti-Mouse NKp46 (CD335) | Thermo Fisher | Cat# 64-3351-80 |
| BV711 Mouse Anti-Human CD5 | BD Biosciences | Cat# 563170 |
| BV785 Mouse Anti-Human CD337 (NKp30) | Biolegend | Cat# 325229 |
| Spark Blue 550 Mouse Anti-Human CD3 | Biolegend | Cat# 344851 |
| PE Mouse Anti-Human CD25 | Thermo Fisher | Cat# 12-0257-41 |
| PE-Cy7 Mouse Anti-Human CD336 (NKp44) | Biolegend | Cat# 325115 |
| AF647 Mouse Anti-Human CD1a | Biolegend | Cat# 300116 |
| APC-H7 Mouse anti-Human CD45 | BD Biosciences | Cat# 560274 |
| APC-Fire810 Mouse anti-Human CD4 | Biolegend | Cat# 344661 |
| Human BD Fc Block | BD Biosciences | Cat# 564220 |
| PerCP/Cyanine5.5 Mouse Anti-Human CD24 | Biolegend | Cat# 311115 |
| BUV615 Mouse Anti-Human CD62L | BD Biosciences | Cat# 751364 |
| PE/Dazzle™ 594 Mouse Anti-Human HLA-A,B,C | Biolegend | Cat# 311439 |
| PE-Cy7 Mouse Anti-Human CCR7 (CD197) | BD Biosciences | Cat# 567314 |
| Ultra-LEAF™ Purified anti-human CD3 Antibody | Biolegend | Cat# 300438 |
| **Chemicals, Peptides, and Recombinant Proteins** | | |
| Recombinant Human BMP-4 Protein | R&D Systems | Cat# 314-BP |
| Recombinant Human VEGF 165 Protein | R&D Systems | Cat#293-VE |
| Recombinant Human Wnt-3a Protein | R&D Systems | Cat#5036-WN |
| Recombinant Human FGF basic/FGF2 (146 aa) Protein | R&D Systems | Cat#233-FB |
| CHIR99021 | Reprocell | Cat#04-0004 |
| Recombinant Human SCF Protein | R&D Systems | Cat#255-SC |
| Recombinant Human Sonic Hedgehog (Shh) | Peprotech | Cat#100-45 |
| Recombinant Human FLT3L | Bio X Cell | No longer made |

| | | |
|---|---|---|
| Recombinant Human TPO | R&D Systems | Cat#288-TP |
| Recombinant Human IL-6 Protein, CF | R&D Systems | Cat#206-IL CF |
| Recombinant Human IL-3 Protein | R&D Systems | Cat#203-IL |
| Recombinant Human IL-11 Protein | Thermo Fisher | Cat#200-11 |
| Recombinant Human IGF-I/IGF-1 Protein, CF | R&D Systems | Cat#291-G1 |
| Human Transferrin | Sigma-Aldrich | Cat#10652202001 |
| Recombinant Human IL-7 Protein | R&D Systems | Cat#207-IL |
| hESC Matrigel Matrix | Corning | Cat#354277 |
| Matrigel Matrix | Corning | Cat#354234 |
| Stemolecule Y27632 | Reprocell | Cat#04-0012-02 |
| Ascorbic Acid | Sigma | Cat#A4544 |
| mTeSR™1 Media | StemCell Technologies | Cat#85850 |
| mTeSR™ Plus Media | StemCell Technologies | Cat#100-0276 |
| StemSpan™ Lymphoid Differentiation Coating Material (100X) | StemCell Technologies | Cat# 09925 |
| StemSpan™ SFEM II | StemCell Technologies | Cat# 09655 |
| StemSpan™ Lymphoid Progenitor Expansion Supplement (10X) | StemCell Technologies | Cat# 09915 |
| StemSpan™ T Cell Progenitor Maturation Supplement | StemCell Technologies | Cat# 09930 |
| X-VIVOTM 15 Serum-free Hematopoietic Cell Medium | Lonza | Cat# 02-053Q |
| StemPro®-34 SFM | Gibco | Cat#10640-019 |
| GlutaMax™ (100x) | Gibco | Cat#35050-061 |
| 1-thioglycerol | Sigma | Cat#M6145 |
| ReLeSR™ | StemCell Technologies | Cat#05873 |
| Primocin | InvivoGen | Ant-pm-2 |
| Doxycycline | Sigma | Cat# D3072 |
| FVS780 | BD Biosciences | Cat# 565388 |

**Table 3.** Antibodies and growth factors used through this study
